# Tumor-initiating genetics and therapy drive divergent molecular evolution in IDH-mutant gliomas

**DOI:** 10.1101/2025.07.11.664189

**Authors:** Tamrin Chowdhury, C. Mircea S. Tesileanu, Emre Kocakavuk, Kevin C Johnson, Jooho Lee, Zeynep Erson-Omay, Chaewon Heo, Kenneth Aldape, Samirkumar B Amin, Kevin J Anderson, David M. Ashley, Jill S. Barnholtz-Sloan, Daniel J. Brat, Andrew R Brodbelt, Ana Valeria Castro, Elizabeth B Claus, Jennifer M. Connelly, Joseph Costello, Indrani Datta, Carol Elliott, Gaetano Finocchiaro, Pim J French, Hui K Gan, Luciano Garofano, Padmaja L Ghospurkar, Anna Golebiewska, Pranav S Gundla, Beth Hermes, Chibo Hong, Youri Hoogstrate, Craig Horbinski, Jason T Huse, Antonio Iavarone, Cihat Karadag, Mustafa Khasraw, Mathilde CM Kouwenhoven, Peter S. LaViolette, Kay Li, Allison Lowman, Katy McCortney, Hyo-Eun Moon, Sanai Nader, MacLean P. Nasrallah, H.K. Ng, D. Ryan Ormond, Marta Padovan, Sun Ha Paek, Laila M Poisson, Sushant Puri, Erica Shen, Mehta Shwetal, Andrew E. Sloan, Wies R Vallentgoed, Erwin G Van Meir, Rachael Vaubel, Taylor Wade, Anna M. Walenkamp, Colin Watts, Tobias Weiss, Micheal Weller, Peter Wesseling, Kerryn Westcott, Bart Westerman, Gordon Y Ye, W. K. Alfred Yung, The GLASS Consortium, Frederick S Varn, Roel GW Verhaak

**Author notes:** These authors contributed equally (Tamrin Chowdhury, C. Mircea S. Tesileanu). These authors jointly supervised the work (Frederick S Varn, Roel GW Verhaak).

## Abstract

Astrocytomas and oligodendrogliomas are slow-growing and treatment-sensitive IDH-mutant gliomas diagnosed at ages 30-50. Local tumor regrowth and treatment resistance is inevitable resulting in 3-10 year astrocytoma and up to >20 years oligodendroglioma survival. We sought to identify genetic changes associated with tumor evolution in response to therapy through multi-timepoint whole-genome/whole-exome sequencing of 206 IDH-mutant glioma patient samples collected through the Glioma Longitudinal Analysis (GLASS) Consortium. We validated known genomic markers of tumor progression, including hypermutation and *CDKN2A* homozygous deletion, and discovered novel genetic alterations that distinguish the response to treatment in astrocytomas compared to oligodendrogliomas. Point mutations in *PIK3CA*, *PIK3R1*, and *NOTCH1* were newly acquired in recurrent oligodendrogliomas and associated with increased mutation rates. Focal oncogene amplifications, together with CDKN2A homozygous deletions, were associated with an increase in recurrence-specific chromosomal imbalances in astrocytomas. Mutational signature analysis revealed additional differences and detected enrichment for the SBS11, and SBS119 mutational signatures after temozolomide treatment in both IDH-glioma subtypes, whereas astrocytomas showed increased ID8 signatures after radiotherapy. These signatures suggest that the genomes of oligodendroglioma and astrocytoma adapt to the selective pressures of tumor progression and treatment in different ways. However, in both IDH-mutant glioma subtypes we observed a convergence of acquired driver gene alterations with genome-wide changes and worse patient outcomes, signaling selection of treatment-refractory clones. By identifying new prognostic markers and delineating the genomic divergence of oligodendrogliomas and astrocytomas after diagnosis, our results suggest that different DNA damage response mechanisms are engaged following chemo- and radiation therapy.

---

Isocitrate dehydrogenase (IDH)-mutant gliomas, named for their gain-of-function mutations in the metabolic enzyme genes *IDH1* and *IDH2*, are the most common primary malignant brain tumors in adults under the age of 50^1–3^. The 2021 WHO classification for Central Nervous System tumors further stratifies IDH-mutant gliomas into two clinical and molecularly distinct subtypes: 1) IDH-mutant oligodendrogliomas (WHO grades 2-3), which present with combined loss of chromosome arms 1p and 19q; and 2) IDH-mutant astrocytomas (WHO grades 2-4), which lack 1p/19q-alterations. Oligodendrogliomas are further marked by mutations in the *TERT* promoter, while IDH-mutant astrocytomas instead prominently feature mutations in *TP53, ATRX*^4,5^. The primary treatment for patients with an IDH-mutant glioma typically involves surgical debulking, followed by alkylating chemotherapy and ionizing radiation while less aggressive tumors (WHO grade 2) may only undergo surgical debulking. More recently, the oral dual IDH1/2 inhibitor vorasidenib has been introduced as a treatment option for selected patients. Despite these therapies, virtually all IDH-mutant gliomas develop treatment resistance, which results in tumor recurrence. IDH-mutant gliomas are diagnosed in the prime of the patient’s lives (median age of diagnosis in the US: oligodendrogliomas = 45 years, and astrocytomas = 37 years)^1^, and ubiquitous emergence of treatment resistance results in a disproportionate number of life-years lost^6^. Some features related to treatment response have been reported, such as alkylating-agent induced hypermutation^7^ and radiation-associated *CDKN2A* homozygous deletions^8^, yet is it currently unknown whether IDH-mutant glioma subtype-specific treatment adaptations exist.

Glioma-specific mutations in *IDH1/2* were first described in 2008 and the 2024 FDA approval of IDH-inhibitor vorasidenib for treatment-naïve progressive WHO grade 2 IDH-mutant gliomas highlights how a deeper understanding of glioma biology can substantially impact development of new therapies^9–11^. While genomic biomarkers or resistance markers for vorasidenib response in IDH-mutant glioma have not yet emerged, IDH inhibition in IDH-mutant leukemias has shown variable treatment responses and resistance has been associated with changes in stemness and mutations in receptor tyrosine kinase genes^12^. Understanding treatment adaptations to chemotherapy, radiotherapy and eventually, IDH-inhibition will provide important insights into how to best sequence or combine therapies.

The Glioma Longitudinal AnalySiS (GLASS) Consortium^13^ seeks to advance our understanding of glioma tumor evolution and response to treatment through genomic characterization of glioma tumors over time^14^. Longitudinal profiling studies from GLASS and others based on exome sequencing datasets on limited (<100) patient numbers have previously demonstrated that a subset of IDH-mutant gliomas treated with alkylating agents acquires a hypermutator genotype^15,16^, that *MYC* gains are associated with shorter post-progression survival^17^ and have shown that radiotherapy increases the frequency of small deletions, homozygous loss of *CDKN2A* and levels of aneuploidy^8,18^. These mutational footprints that drive tumor progression and therapeutic resistance were either pre-existent in small tumor cell populations and selected for or are the direct result from the DNA damage response triggered by radiation and alkylating chemotherapy, and providing a rationale for targeting the engaged biological networks^19^. In this study, we characterize the genetic responses to treatment of IDH-mutant glioma through analysis of whole-genome and whole-exome sequencing multi-timepoint datasets from 206 patients with highly detailed clinical annotation included in the GLASS Data Resource^14^. Our findings demonstrate the extent to which treatment shapes driver gene alterations, mutational signatures, chromosomal aneuploidy, and tumor clonality, revealing the distinct evolutionary trajectories across IDH-mutant gliomas with and without 1p/19q-codeletion.

## Clinical characteristics and patient timelines

Tumor specimens and clinical annotation from patients diagnosed with an IDH-mutant glioma were contributed by 21 institutions across 10 countries, including Australia, Brazil, China, France, Germany, Japan, Luxembourg, South-Korea, the United Kingdom, and the United States (**Fig. 1A**). Due to the longitudinal nature of our study, patients had undergone at least two time-separated surgical procedures to acquire tumor tissue, which were typically debulking resections (92%) and biopsies for the remaining cases and timepoints. Longitudinal sequencing datasets were generated on specimens from 238 patients. After applying quality filters, tumor samples from 206 patients with mutational profiles and tumor samples with mutational as well as DNA copy number profiles from 183 patients at two timepoints were retained for downstream analyses, selecting the first and last available tumor samples in cases of multiples (**Fig. 1B**). Across the cohort, 145 tumor pairs were obtained at the time of diagnosis (Tx) and first recurrence (R1), while 61 tumor pairs were included with other surgery timing combinations such as Tx-R2 (20 tumor pairs), and R1-R2 (29 tumor pairs). Analyses reflecting results from all tumor pairs are referred to as initial-recurrence, analyses including only primary-recurrent pairs are indicated accordingly.

**Figure 1.**
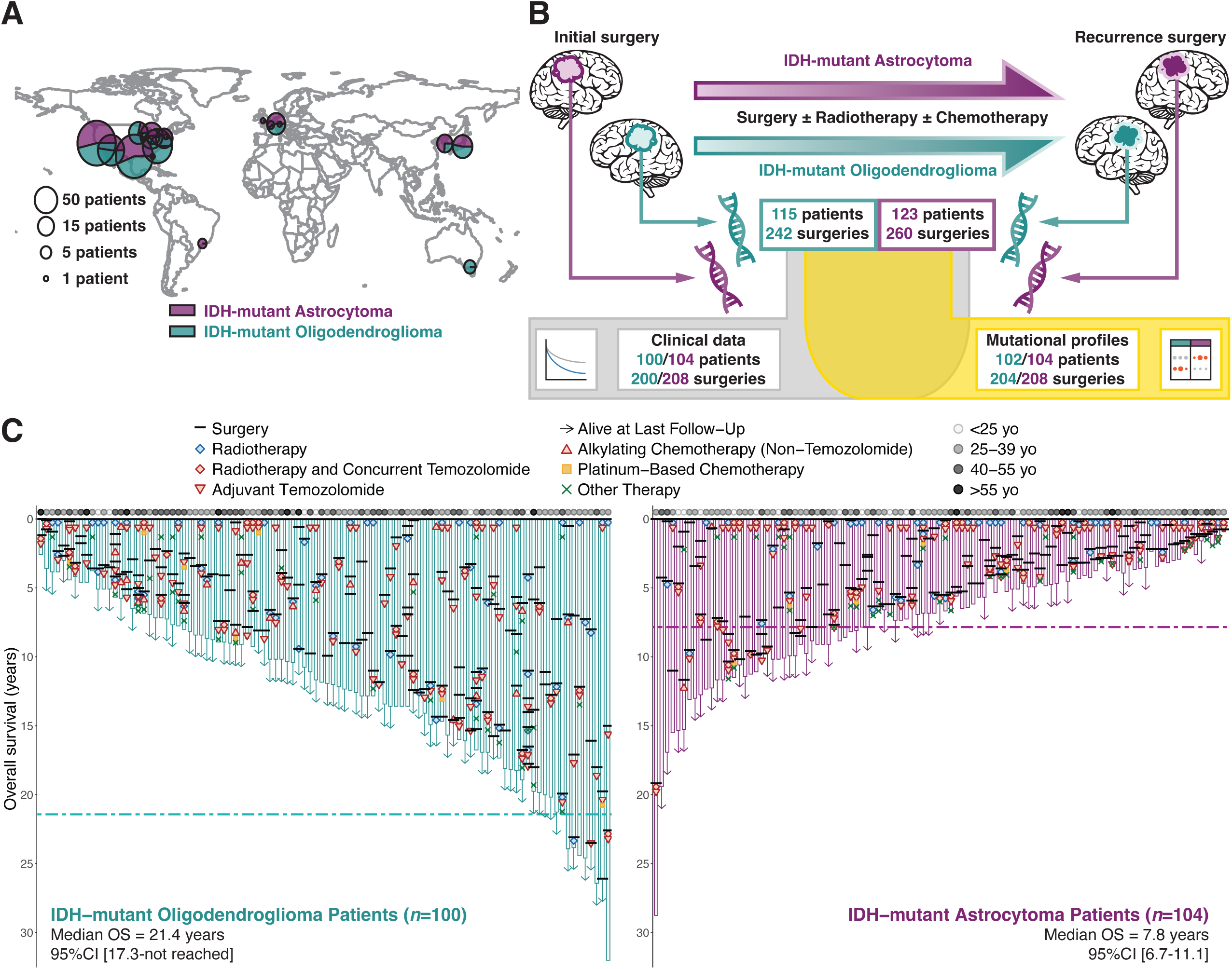
GLASS: a comprehensive global dataset on IDH-mutant glioma evolution. **A.** The GLASS cohort represents patients from five continents as shown in the number of patients per contributing city and the proportions of the individual glioma types. **B.** The GLASS IDH-mutant cohort includes 238 patients with an IDH-mutant glioma from whom longitudinal molecular and/or clinical data were collected across at least two time-separated surgeries. **C.** Extensive clinical data have been acquired with respect to treatment and prognostic factors; overall survival is shown for 204 patients of which 110 patients (70 with oligodendrogliomas and 41 with astrocytomas) were still alive at last follow-up as specified by the arrows. As expected, the median overall survival per glioma type (dotted lines) differed significantly in favor of the oligodendroglioma cohort. Abbreviations: OS, overall survival; CI, confidence interval.

The extent of clinical follow-up for the GLASS cohort was a strength of our study. Clinical annotation was collected for all patients and terminology was standardized across institutions. In addition to patient demographics, tumor pathology (2021 WHO grade, histology), surgery details (extent of resection versus biopsy, tumor location), treatment regimens (chemotherapy, radiation therapy, other treatments) and treatment outcomes (surgical interval, overall survival) were obtained. Patients from our cohort were diagnosed at a significantly younger age compared to patients with IDH-mutant glioma from The Cancer Genome Atlas (TCGA) (Wilcoxon test *p*-value 0.013 and <0.001 for astrocytomas and oligodendrogliomas, respectively; **Extended Data Fig. 1A**)^20^. GLASS oligodendroglioma patients showed longer overall survival from the time of diagnosis compared to TCGA, while overall survival from the time of birth was significantly shorter for both GLASS oligodendrogliomas and astrocytomas (**Extended Data Fig. 1B and 1C**). These cohort differences may have resulted from an inclusion bias towards patients who have undergone multiple surgeries.

The median follow-up for the GLASS cohort was 6.5 years for astrocytoma patients and 11.9 years for oligodendroglioma patients (**Table 1**), compared to 2.3 years and 1.9 years, respectively, in TCGA (Wilcoxon: *p* < 0.001 for both). Tumor grade increased between timepoints in 59% of astrocytoma and 47% of oligodendrogliomas. Throughout their treatment, 76% of GLASS IDH-mutant glioma patients received alkylating chemotherapy, with temozolomide as the preferred alkylating agent administered (astrocytomas: 74%, oligodendrogliomas: 70%)(**Fig. 1C**). Platinum-based chemotherapy was the second chemotherapy of choice and was always used in addition to alkylating chemotherapy (astrocytomas: *n* = 6, oligodendrogliomas: *n* = 7). Radiotherapy was used to treat 72% of astrocytoma patients and 69% of oligodendroglioma patients. Other modes of therapy were attempted in 32% of IDH-mutant glioma patients and included treatment strategies such as anti-VEGF therapy (*n* = 35) and retinol-based drugs such as all-trans retinoic acid (*n* = 5). A small number of patients (astrocytomas: *n* = 16, oligodendrogliomas: *n* = 14) were not reported to receive any treatment during their known follow-up trajectories. At time of diagnosis, primary treatment with radiotherapy was significantly associated with poor overall survival (*p* = 0.005 and *p* < 0.001, log-rank test) but not time to secondary surgery (*p* = 0.48 and *p* = 0.24, log-rank test) in both astrocytoma and oligodendroglioma patients, reflecting selection bias to more aggressively interview in patients with an IDH-mutant glioma with perceived poor prognosis (**Extended Data Table 1 to 4**). Higher tumor grade but not patient age was significantly associated with patients received primary radiation treatment. A minor subset (23%) of the cohort received both radiotherapy and alkylating agents as primary treatment.

**Table 1.** Clinical characteristics of patients with IDH-mutant glioma from GLASS cohort at baseline. Categorical variables were compared using Fisher’s exact test, and continuous variables with the Wilcoxon rank-sum test or the Kruskal–Wallis test, as appropriate. Abbreviations: Min, minimum, Max, maximum.

|  |  | IDH-mutant Astrocytoma<br>N = 104 | IDH-mutant Oligodendroglioma<br>N = 100 | p-value |
| --- | --- | --- | --- | --- |
| **Demographics** |  |  |  |  |
| *Age at diagnosis (years)* |  |  |  | 0.005 |
|  | Median [Min, Max] | 35 [14, 67] | 37 [17, 70] |  |
| *Sex* |  |  |  | >0.9 |
|  | Female | 38% (39/104) | 38% (39/100) |  |
|  | Male | 63% (65/104) | 62% (62/100) |  |
| *Follow-up (years)* |  |  |  | <0.001 |
|  | Median [Min, Max] | 6.5 [0.9, 28.8] | 11.9 [2.6, 32.0] |  |
| **Primary Treatment** |  |  |  |  |
| *Surgery* |  |  |  | 0.4 |
|  | Biopsy | 8.8% (5/57) | 13% (10/77) |  |
|  | Resection | 91% (52/57) | 87% (67/77) |  |
|  | Unknown | 47 | 23 |  |
| *Radiotherapy* |  |  |  | 0.015 |
|  | Non-treated | 51% (48/95) | 69% (55/80) |  |
|  | Treated | 49% (47/95) | 31% (25/80) |  |
|  | Unknown | 9 | 20 |  |
| *Alkylating chemotherapy* |  |  |  | 0.10 |
|  | Non-treated | 53% (46/87) | 65% (51/78) |  |
|  | Treated | 47% (41/87) | 35% (27/78) |  |
|  | Unknown | 17 | 22 |  |
| *Platinum-based chemotherapy* |  |  |  | 0.5 |
|  | Non-treated | 100% (22/22) | 93% (28/30) |  |
|  | Treated | 0% (0/22) | 6.7% (2/30) |  |
|  | Unknown | 82 | 70 |  |
| *Other therapies* |  |  |  | 0.2 |
|  | Non-treated | 59% (13/22) | 77% (23/30) |  |
|  | Treated | 41% (9/22) | 23% (7/30) |  |
|  | Unknown | 82 | 70 |  |
| **Primary Pathology** |  |  |  |  |
| *Grade* |  |  |  | 0.002 |
|  | 2 | 73% (66/91) | 92% (67/73) |  |
|  | 3 | 14% (13/91) | 8.2% (6/73) |  |
|  | 4 | 13% (12/91) |  |  |
|  | Unknown | 13 | 27 |  |

Samples were analyzed using whole-genome sequencing (astrocytomas: 111 samples, 53 patients; oligodendrogliomas: 178 samples, 84 patients), whole-exome sequencing (astrocytomas: 203 samples, 97 patients; oligodendrogliomas: 218 samples, 103 patients), and bulk RNA sequencing (astrocytomas: 95 samples, 50 patients; oligodendrogliomas: 112 samples, 57 patients)(**Fig. 1A**). After rigorous quality checks, a clinical annotation dataset (astrocytomas: 208 samples, 104 patients; oligodendrogliomas: 200 samples, 100 patients) was established for which longitudinal sequencing datasets were available in a majority subset (astrocytomas: 188 samples, 94 patients; oligodendrogliomas: 178 samples, 89 patients), and mutational profiles, DNA copy number alterations and mutational signatures were determined. All processed genomic data and clinical annotations are available via https://www.synapse.org/glass.

## IDH-mutant oligodendrogliomas and astrocytomas acquire different genomic driver events

While IDH-mutant gliomas generally respond well to chemoradiation, disease recurrence is inevitable^21^. We previously reported that a subset of IDH-mutant gliomas acquire genomic alterations such as somatic hypermutation or *CDKN2A* homozygous deletion at the time of disease progression, which were associated with alkylating chemotherapy and radiotherapy, respectively^7,8,22^. Leveraging the larger GLASS cohort provided power to detect new drivers of tumor progression by comparing significant gene-level events at the time of diagnosis and disease recurrence. At diagnosis, we found significant enrichment of mutations in canonical driver genes (astrocytomas: *TP53* and *ATRX*; oligodendrogliomas: *TERT* promoter mutations, *CIC* and *FUBP1*), which were preserved at recurrence (**Fig. 2A**)^4^. As expected, a subset of both astrocytomas (29%) and oligodendrogliomas (23%) that received alkylating chemotherapy recurred with a significantly higher number of mutations (> 10 mut/Mb) and were labeled as hypermutators. Amongst non-hypermutated recurrent oligodendrogliomas (*n* = 70), we newly discovered a significant enrichment in *NOTCH1*, *PIK3CA* and *PIK3R1* (**Fig. 2B**). *NOTCH1* and *PIK3CA/PIK3R1* mutations found at recurrence were not associated with specific prior treatments and showed mutual exclusivity (DISCOVER algorithm^23^: *p* = 0.046 at primary, and *p* = 0.036 at non-hypermutant recurrence)(**Extended Data Fig. 2A**). In leukemias, NOTCH1 has been implicated as a regulator of PTEN with downstream effects on PI3K signaling which may explain why in oligodendrogliomas PI3K and NOTCH1 mutations are mutually exclusive^24,25^. *NOTCH1* mutations in oligodendrogliomas have been associated with reduced cellular differentiation and mouse models have shown that models harboring *Notch1* mutations are resistant to the cell lineage differentiation effects of mutant IDH inhibitors, nominating *NOTCH1* mutations a potential IDH-inhibitor resistance biomarker in oligodendrogliomas^26,27^. We observed the previously reported significant enrichment in *MSH6* mutations at recurrence as well as newly acquired *PIK3CA* mutations in hypermutated oligodendrogliomas (mutations at diagnosis: *MSH6* = 0/10, *PIK3CA* = 2/10; mutations at recurrence: *MSH6* = 12/16, *PIK3CA* = 13/16). We observed a modest and non-significant increase in hypermutated IDH-mutant gliomas with MSH6 mutation compared to MSH6 wildtype hypermutants, a consistent trend in both GLASS and the AACR GENIE cohort^28^(**Extended Data Fig. 2B**). While some hypermutated astrocytomas acquired *MSH6* mutations (*n* = 7/20), this enrichment did not reach statistical significance. Consistent with this, the *MSH6* mutation frequency was significantly lower in astrocytomas compared to oligodendrogliomas (Fisher’s exact test, *p* = 0.048)(**Fig. 2B**)^7^. In contrast to oligodendroglioma, no gene mutations were significantly associated with recurrence in astrocytomas regardless of hypermutation status.

**Figure 2.**
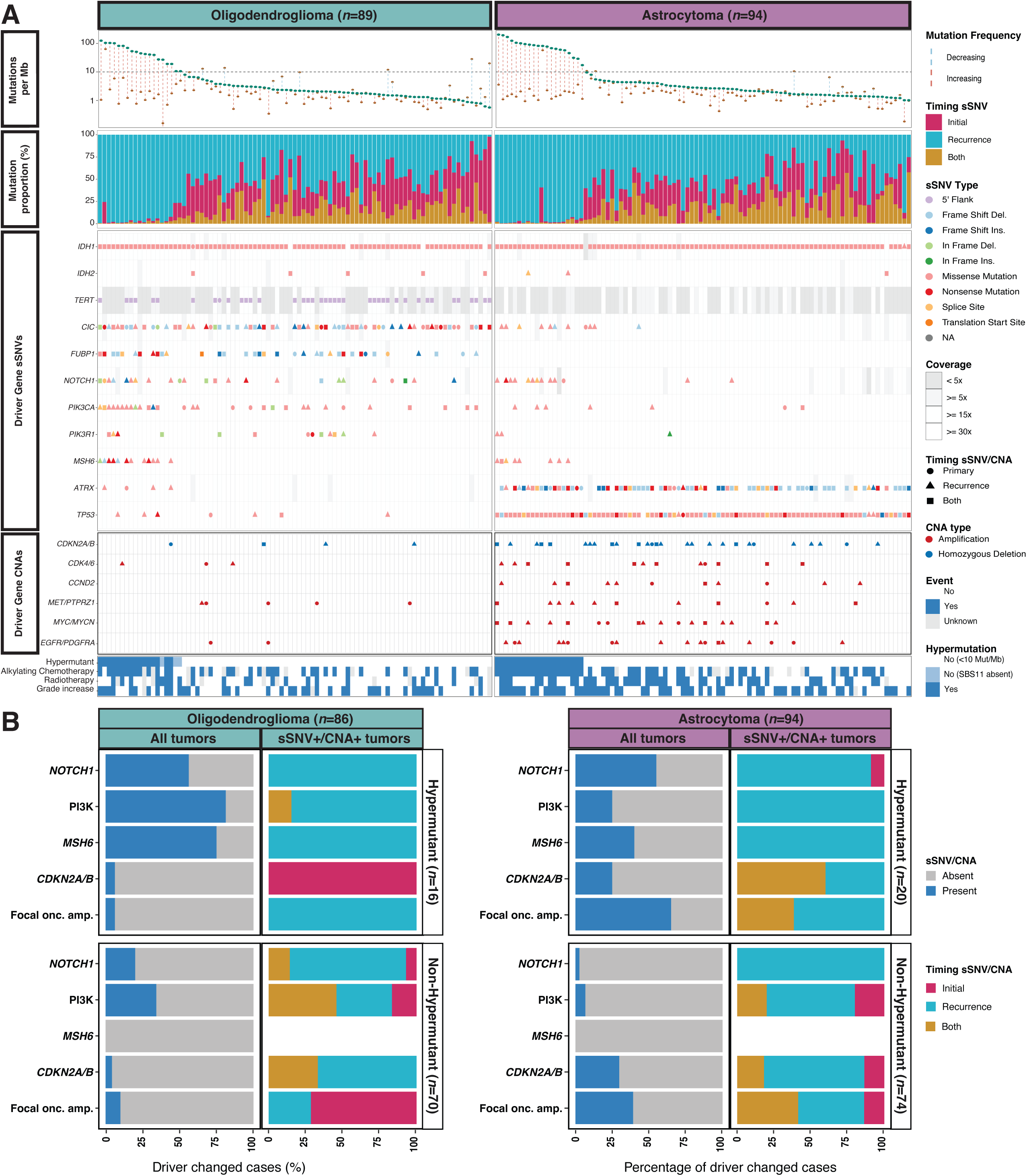
sSNVs drive oligodendroglioma progression whereas astrocytomas progress through CNAs. **A.** sSNV/CNA landscape of patients with mutation and DNA copy number profiles from at least two timepoints (*n*=183) in which individual patients (columns) are ordered by mutation frequency at recurrence. Segments ordered top to bottom: i. Mutation frequency increased over time in 81% of gliomas and decreased in the remaining 19%. At recurrence, 39 gliomas surpassed the sSNV threshold for hypermutation (10 Mut/Mb, black dotted line); ii. Gliomas with the highest mutation frequency at recurrence (left) had mostly recurrence-only mutations, whereas those with lower mutation frequencies (right) were comprised of mainly primary-only and timing-shared mutations; iii. Oligodendrogliomas had canonical mutations in *CIC*, *FUBP1*, and the *TERT* promoter; astrocytomas in *ATRX* and *TP53*. Non-hypermutant oligodendrogliomas were significantly enriched for *NOTCH1*, *PIK3CA*, and *PIK3R1* mutations in dN/dS and MutSigCV analyses. Temozolomide-induced hypermutant gliomas showed broad sSNV driver enrichment, possibly linked to recurrence-specific *MSH6* mutations; iv. Focal oncogene amplifications (*CDK4*, *CDK6*, *CCND2*, *MET*, *PTPRZ1*, *MYC*, *MYCN*, *EGFR*, and *PDGFRA*) and homozygous deletions of *CDKN2A/B* were scarce in recurrent oligodendrogliomas, but they were significantly increased in recurrent astrocytomas; v. Clinical markers to relate to panels above. **B.** Highlights of landscape (panel **A.**iii and iv) excluding those without SBS11 expression. Left panels indicate presence of sSNV/CNA events, right panels represent sSNV/CNA timing. Abbreviations: Mb, megabase; sSNV, somatic single nucleotide variant; CNA, copy number alteration; Mut, mutations; SBS11, single-base substitutions signature 11; PI3K, *PIK3CA* and *PIK3R1* combined; Onc. amp., oncogene amplifications.

We used the GISTIC algorithm to identify focal DNA copy number gains and losses that target well-described oncogenes and tumor suppressor genes, focusing on events detected at recurrence. Focal *CDKN2A/B* homozygous deletions were acquired in recurrent astrocytomas with statistically significant enrichment (McNemar’s test: *p* = 0.001) but not oligodendrogliomas (McNemar’s test: *p* = 0.48). Fifteen of 19 gliomas with acquired *CDKN2A/B* homozygous deletion were pre-treated with radiotherapy suggesting that radiation provides a selective pressure favoring CDKN2A-inactive cells (odds ratio = 2.71, *p*-value = 0.01, Fisher’s exact test). We newly report that focal high-level oncogene amplifications were detected at significantly increased frequency in recurrent astrocytomas. In total, 38 of 94 recurrent astrocytomas contained at least one focal oncogenic amplification, compared to 19 of 94 initial astrocytomas. Frequently amplified oncogenes included *MET*/*PTPRZ1*, two oncogenes that are located relatively adjacent to each other on 7q31, *MYC* (8p24), *CDK4* (12q14), *CCND2* (12p13), *EGFR* (7p11), and *PDGFRA* (4q12). Related to these findings, *MET-PTPRZ1* transcript fusions have recently been reported in glioma and found to be responsive to targeted *MET* inhibitors (**Fig. 2A and B**)^29–31^. In contrast, there was no significant enrichment of amplifications or homozygous deletions in recurrent oligodendrogliomas.

Hypermutation at recurrence has been shown to be an unfavorable prognostic factor for both IDH-mutant glioma subtypes^7,32^ and this correlation was confirmed in our cohort (**Fig. 3A and 3B**). Enrichment of PI3K and *NOTCH1* mutations did not further stratify survival in non-hypermutant tumors in GLASS (**Fig. 3C**). However, PI3K/*NOTCH1*-mutant oligodendrogliomas did associate with worse overall survival in the AACR GENIE validation set (**Fig. 3D; Extended Data Fig. 2D**). Homozygous deletions of *CDKN2A/B* at astrocytoma recurrence associated with poor patient overall survival (median: 1.1 years vs 3.6 years), confirming the use of *CDKN2A/B* loss as a grade 4 marker in the 2021 WHO classification (**Extended Data Fig. 2C, Extended Data Table 5 and Extended Data Table 6**). Acquired focal amplification of at least one well-described oncogene (*MET/PTPRZ1/CDK4/CCND2/MYC/MYCN/EGFR/PDGFRA*) also associated with significantly shorter overall survival in patients with non-hypermutated astrocytomas only (median: 2.1 years vs 4.6 years)(**Extended Data Fig. 2C**). When combined, acquired *CDKN2A/B* homozygous deletion and focal oncogene amplification represented a significant molecular marker designating inferior patient outcome (**Fig. 3E**). Multivariable analysis was unable to show that the effect of the individual markers was independent, possibly due to the low sample size. However, the presence of *CDKN2A* homozygous deletions or of focal oncogene amplifications in AACR GENIE astrocytoma patients were independently associated with worse outcomes (**Fig. 3F; Extended Data Fig. 2E**). In summary, we found that oligodendrogliomas showed enrichment for acquired canonical mutations in driver genes at recurrence, whereas astrocytoma disease progression was associated with significant enrichment in acquired focal DNA copy number alterations.

**Figure 3.**
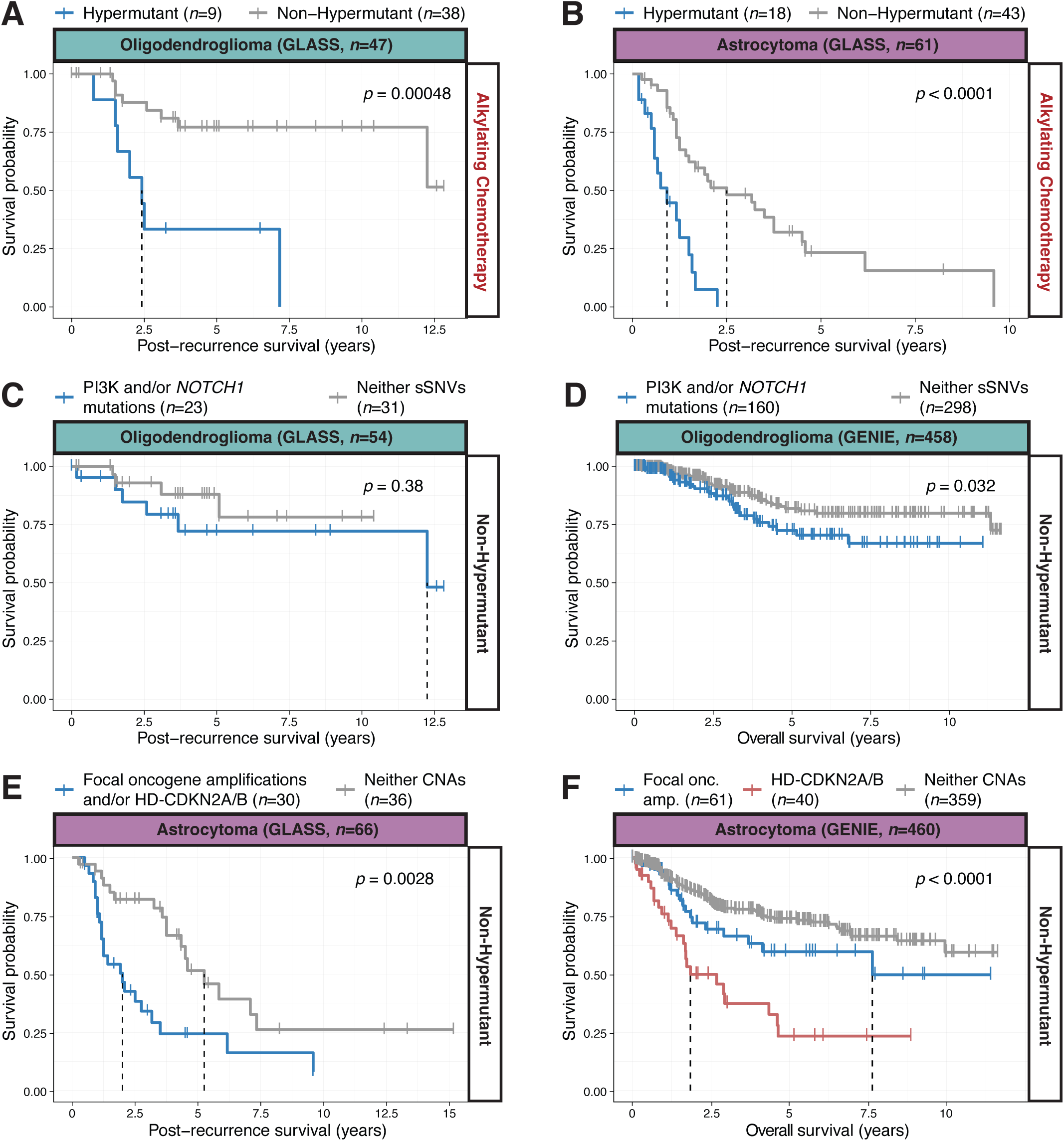
Post-recurrence survival is associated with acquired hypermutation and CNAs. Kaplan-Meier curves comparing post-recurrence survival of different patient groups (GLASS cohort **A-C** and **E**; GENIE cohort **D** and **F**). Statistical significance assessed using log-rank tests. GLASS cohort patients who received alkylating chemotherapy prior to recurrence to treat (**A**) oligodendrogliomas and (**B**) astrocytomas; patients with gliomas which underwent hypermutation transformation associated with worse post-recurrence survival than those without hypermutation transformation. **C.** Survival analysis of GLASS cohort oligodendroglioma patients with acquired PI3K or *NOTCH1* mutations compared to patients whose tumors did not acquire such events. **D.** In GENIE cohort patients, PI3K and *NOTCH1* mutations were correlated with worse overall survival in non-hypermutant oligodendrogliomas. **E.** Post-recurrence survival of GLASS cohort patients with a non-hypermutant astrocytoma and a focal oncogene amplification and/or homozygously deleted *CDKN2A/B* compared to patients with a non-hypermutant astrocytoma lacking those variants. **F.** Both homozygous deletion of *CDKN2A/B* (±focal oncogene amplifications) and focal oncogene amplifications only (i.e. without HD-CDKN2A/B) were correlated with worse overall survival in GENIE cohort patients with non-hypermutant astrocytomas. Abbreviations: PI3K, *PIK3CA* and *PIK3R1* combined; sSNVs, somatic single nucleotide variants; Focal oncogene, *CDK4*, *CDK6*, *CCND2*, *MET*, *PTPRZ1*, *MYC*, *MYCN*, *EGFR*, or *PDGFRA*; HD-CDKN2A/B, homozygous deletion of *CDKN2A/B*; CNAs, copy number alterations; onc. amp., oncogene amplifications.

## Increased aneuploidy rates in astrocytoma but not oligodendroglioma

Aneuploidy levels have been associated with worse patient outcomes in astrocytoma but not oligodendroglioma^33,34^. We evaluated broad DNA copy number alterations patterns in association to treatment status and acquired genomic drivers. Newly diagnosed oligodendrogliomas exhibited few DNA copy number changes beyond 1p/19q-codeletion, whereas astrocytomas harbored higher levels of aneuploidy, defined as broad low-level DNA copy number changes (**Fig. 4A; Extended Data Fig. 3A**)^4^. The level of aneuploidy, defined as the fraction of genome that was copy-number altered, increased with grade (**Extended Data Fig. 3B**). In addition to the canonical 1p/19q-codeletions, our analysis revealed frequent chromosome-arm level loss of 4p and 4q in oligodendrogliomas, further enriched at recurrence (23% to 33%) while gain of 7q and loss of 10q were frequently detected in astrocytomas (32%)(**Fig. 4B, Extended Data Fig. 3C, Extended Data Fig. 3D**). At recurrence, we observed a further significantly increased fraction of genome altered in astrocytomas but not oligodendrogliomas (**Fig. 4C**). The amount of change in the fraction of genome altered and the length of the interval between initial and recurrent astrocytoma were significantly associated (**Extended Data Fig. 3E**). This observation was paired with a high number of shared broad copy number alterations in oligodendrogliomas while most chromosomal gains or losses in astrocytomas were private to either primary or recurrent tumor (**Fig. 4A**).

**Figure 4.**
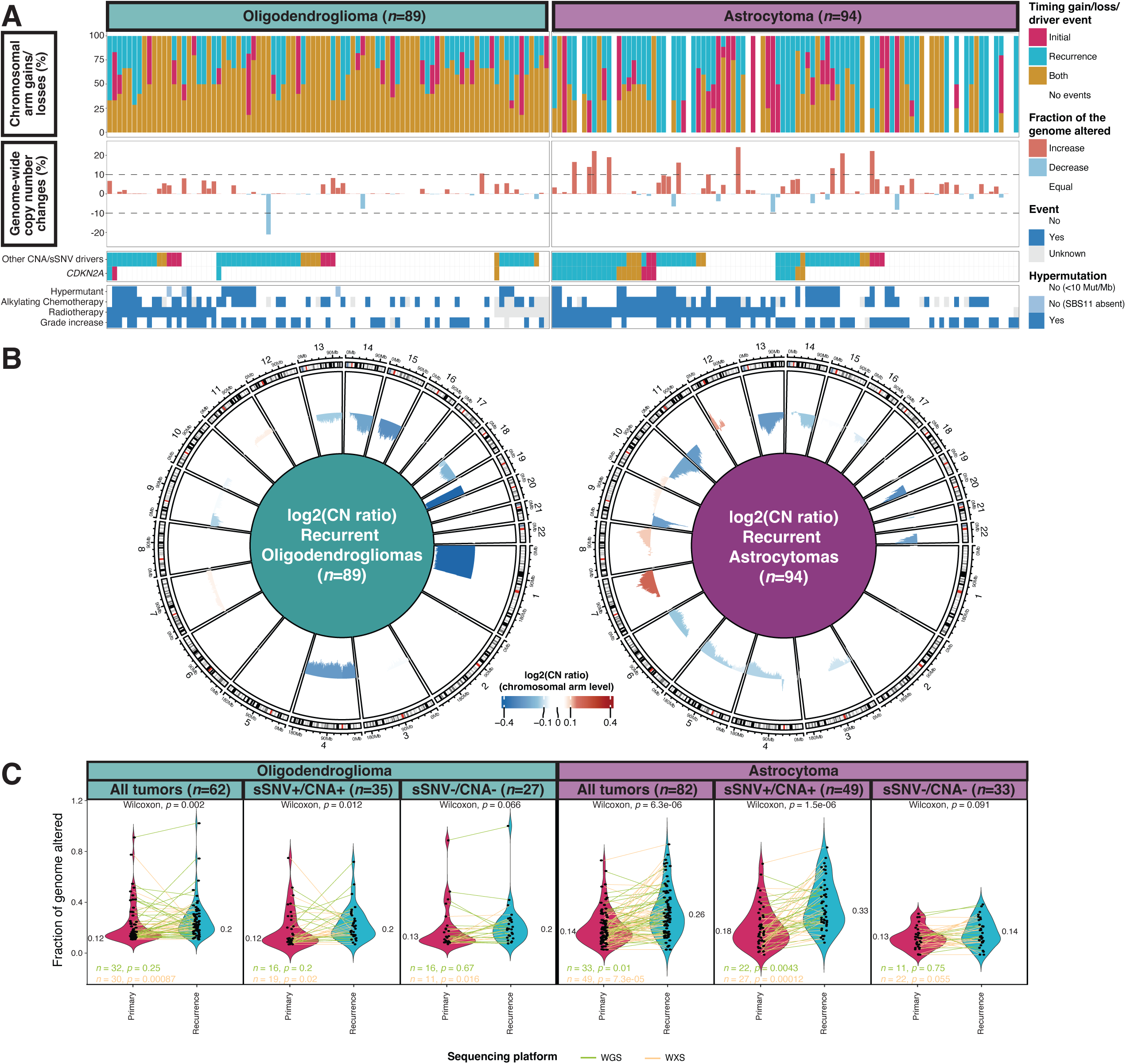
Aneuploidy rises in recurrent astrocytomas but stays relatively stable in oligodendrogliomas. **A.** Aneuploidy landscape of patients with mutation and DNA copy number profiles from at least two timepoints (*n*=183) in which individual patients (columns) are ordered by sSNV/CNA driver status. Recurrent astrocytomas showed higher numbers of recurrent-specific aneuploidies at both arm-level and genome-wide scales compared to oligodendrogliomas. Segments ordered from top to bottom: i. Shared and private copy number segment fractions by tumor; ii. Ratio of the fraction of genome altered between initial and recurrent tumor. Higher value indicates a higher fraction of genome altered in the recurrent tumor; iii. Significant molecular markers to relate to panels above; iv. Clinical markers to relate to panels above. **B.** Recurrent astrocytomas showed a higher number of significant chromosomal arms gains and losses than oligodendrogliomas (mean log2(CN ratio) ≤ −0.1 or ≥ 0.1: recurrent oligodendrogliomas = 10 chromosomal arms; recurrent astrocytomas = 14 chromosomal arms). **C.** Comparison of genome-wide aneuploidy (measured as fraction of genome altered) between primary-only and matching recurrent gliomas, grouped by glioma type and the presence of newly acquired driver genes. The following driver gene alterations were considered: focal oncogene amplifications (*CDK4*, *CDK6*, *CCND2*, *MET*, *PTPRZ1*, *MYC*, *MYCN*, *EGFR*, and *PDGFRA*) and homozygous deletions of *CDKN2A/B*; *NOTCH1*, *PIK3CA*, and *PIK3R1* somatic single-nucleotide variants. Abbreviations: CNA, copy number alteration; sSNV, somatic single nucleotide variant; CN, copy number; Mut, mutations; Mb, megabase; SBS11, single-base substitutions signature 11; WGS, whole-genome sequencing; WXS, whole-exome sequencing.

We found that the significant increase in broad copy number changes was only observed in astrocytomas that acquired a driver gene alteration (*CDKN2A* homozygous deletion or focal oncogene amplification), suggesting that an increased sensitivity to acquired broad copy number alterations may predispose to focal DNA lesions (**Fig. 4C**). High aneuploidy levels have been linked with decreased response to immune checkpoint inhibition^35^ and may also induce therapeutic vulnerabilities to agents such as KIF18A inhibitors^36^. Our findings provide a rationale for targeting aneuploidy in astrocytomas but not oligodendrogliomas.

## Distinct mutational processes shape the oligodendroglioma and astrocytoma genomes

The median whole-genome sequencing derived somatic single base substitutions (SBS) tumor mutation burden (TMB) of primary oligodendrogliomas and astrocytomas was 0.9 mut/Mb and 1 mut/Mb, respectively, which is considered a low mutational burden relative to other tumor types^37^. Mutational burden at diagnosis significantly increased with higher tumor grade (**Extended Data Fig. 4A**). At recurrence, and excluding hypermutators, TMB (comprising SBS, double-base substitutions, small insertions and deletions) significantly increased in both oligodendrogliomas and astrocytomas. However, in oligodendrogliomas, TMB increases were only significantly associated with tumors acquired driver gene alterations (**Fig. 5A**). Mutations shared between timepoints are expected to be mostly truncal, reflecting early derivation in the evolution of a given tumor^15^. Among all mutations detected in the initial tumor, the median fraction that was also detected in the recurrent tumor was 36% for oligodendrogliomas and 45% for astrocytomas (**Extended Data Fig. 4B**). Exome sequencing datasets from different sectors of the same tumor are available for a subset of GLASS (*n* = 5) and the median fraction of spatially shared mutations in IDH-mutant gliomas was 53%, suggesting that spatial heterogeneity accounts for part of the lack of overlap in mutations between initial and recurrent tumors (**Extended Data Fig. 4C**). The relatively low number of spatially and longitudinally shared mutations suggests a relative absence of selection processes, enabling new clones to continuously arise and compete for resources. The length of the surgical interval was negatively correlated with the fraction of mutations shared by primary-recurrent tumor pairs in both IDH-mutant subtypes, and the recurrence-only mutational fraction significantly correlated with surgical interval in oligodendrogliomas (**Fig. 5B**).

**Figure 5.**
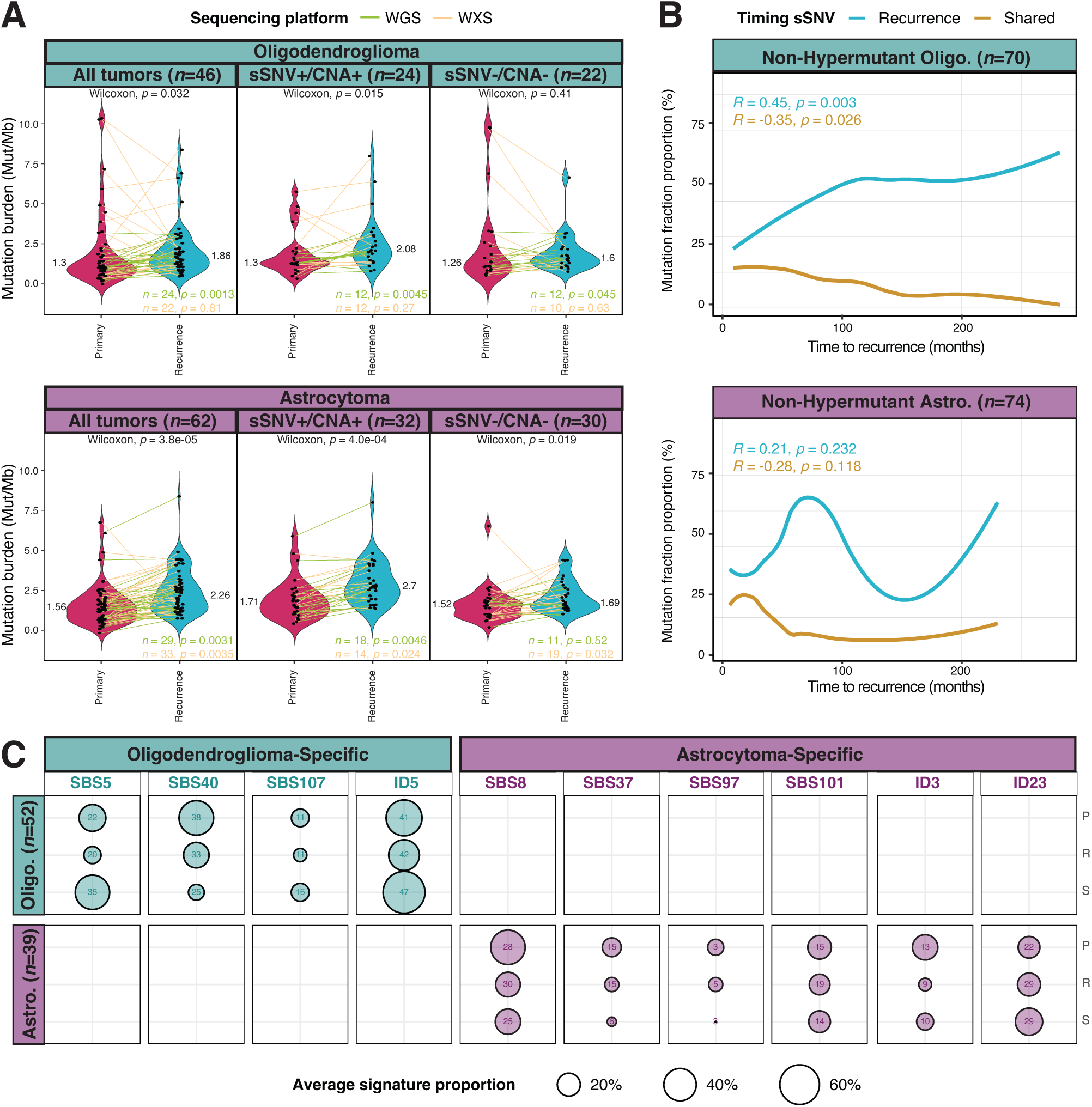
Recurrent gliomas show increased tumor mutational burden and glioma type-specific mutational divergence tied to acquired drivers and surgical interval. **A.** Comparison of mutation burden (number of mutations per megabase) between primary-only and matching recurrent gliomas, grouped by glioma type and the presence of newly acquired driver genes. The following driver gene alterations were considered: focal oncogene amplifications (*CDK4*, *CDK6*, *CCND2*, *MET*, *PTPRZ1*, *MYC*, *MYCN*, *EGFR*, and *PDGFRA*) and homozygous deletions of *CDKN2A/B*; *NOTCH1*, *PIK3CA*, and *PIK3R1* somatic single-nucleotide variants. **B.** In non-hypermutant oligodendrogliomas, recurrence-only mutations showed a significant correlation with time between surgeries while shared mutations declined over time; a pattern which was not present in the astrocytomas. **C.** SBS and ID signatures were compared using the average percentage of alterations attributed to these signatures (circle size) and the number of patients with any alteration attributable to these signatures (number in circle) per glioma type and timing of the signatures. In oligodendrogliomas, 4 treatment-agnostic SBS and ID signatures were detected which were not seen in astrocytomas (Oligodendroglioma-Specific), and conversely 6 other signatures were exclusively detected in astrocytomas (Astrocytoma-Specific). Abbreviations: WGS, whole-genome sequencing; WXS, whole-exome sequencing; sSNV, somatic single nucleotide variant; CNA, copy number alteration; Mut, mutations; Mb, megabase; SBS, single-base substitutions; ID, small insertions and deletions; Astro., astrocytomas; Oligo., oligodendrogliomas; P, primary; R, recurrence; S, shared.

Mutational signatures reflect the underlying mutational processes in a tumor and may be associated with external environmental exposures or internal biological processes such as defective DNA repair. Other than ionizing radiation, no carcinogenic exposures have been causally linked to incidence risk in IDH-mutant glioma^38^. We explored the developmental history of IDH-mutant gliomas via the analysis of mutational signatures from whole-genome sequencing datasets. Mutational signatures were extracted from 78 genomes from 39 astrocytomas and 112 genomes from 56 oligodendrogliomas. This analysis revealed similar signatures to those previously reported signatures in the combined catalogue of the COSMIC and Signal databases^39,40^, including 28 SBS or DBS signatures and 12 ID signatures. Amongst mutations detected in untreated primary tumors or mutations shared between primary and matching recurrent tumors, several SBS (*n* = 6) and ID (*n* = 7) signatures were found in more than five samples, such as the age-related clock-like SBS1, ID1, and ID2 signatures (**Extended Data Fig. 5**). We observed the presence of SBS120 across both IDH-mutant glioma types, an intriguing signature of unknown origin that is unique to tumors in the central nervous system^40^. The SBS5, SBS8 and SBS40 signatures are associated with late replication regions in different genomic contexts and were differentially observed between oligodendrogliomas (SBS5/SBS40) or astrocytomas (SBS8)^41^(**Fig. 5C**). The C>T SBS97 signature has been related to mismatch repair deficiencies and was found only in astrocytomas^42^ together with SBS37 and SBS101, signatures of unknown etiology. In contrast, C to A transversions dominated the SBS107 signature and was detected in oligodendrogliomas but not astrocytomas. This signature shows similarity to the SBS4 tobacco smoking signature. Previous SBS107 associations were found in bladder, renal and liver carcinomas^40,43^. Signatures ID3, linked to exposure to tobacco smoke, and ID23, associated with aristolochic acid exposure^44^, were detected in subsets of astrocytomas. Smoking is not associated with an appreciably elevated risk of IDH-mutant glioma and aristolochic acid exposure additionally results in increased fractions of SBS22a, SBS22b which were not detected, suggesting alternative mechanisms are able to generate SBS107, ID3 and ID23^45^. ID5 is reported as a clock-like signature and was found in oligodendrogliomas but not astrocytomas^39^. We compared mutational signatures in tumors from newly diagnosed patients across different geographical locations and observed a significant enrichment of ID2 mutations in four patients from South-Korea/Hong Kong compared to the rest of the cohort (*p* = 0.025). ID2 mutations are associated with normal replication slippage, and the increased rate in Asian patients may reflect the reported enrichment of glioma-associated risk alleles in *ERCC1*^46^. In summary, we identified significant differences in mutational signatures between oligodendrogliomas and astrocytomas, suggesting that a surprising variety in different mechanisms underlie the initiation, DNA damage repair response and clonal selection processes of both glioma subtypes.

## Therapy-associated mutational signatures

Therapeutic bottlenecks drive clonal selection of tumor cells able to effectively repair treatment-induced DNA damage, and their mutational signatures may represent the footprint of the associated DNA damage repair mechanisms. We evaluated the enrichment of mutational signatures after therapy exposure relative to matched pre-treatment samples. The temozolomide-related hypermutation signature SBS11 was detected in 29% respectively 23% of temozolomide-treated astrocytomas and oligodendrogliomas (**Fig. 6A and 6B**). We discovered a new temozolomide-treated signature in 8 out of 46 non-hypermutated oligodendrogliomas, which significantly acquired SBS119. SBS11 is dominated by C>T transversions, while SBS119 is enriched for T>C substitutions and shows similarity with microsatellite instability associated signatures such as SBS26. The SBS119 signature was also detected in 3 of 33 temozolomide-treated non-hypermutated astrocytomas, while absent in primary tumors and cases that were not treated with temozolomide (**Fig. 6A and 6B**).

**Figure 6.**
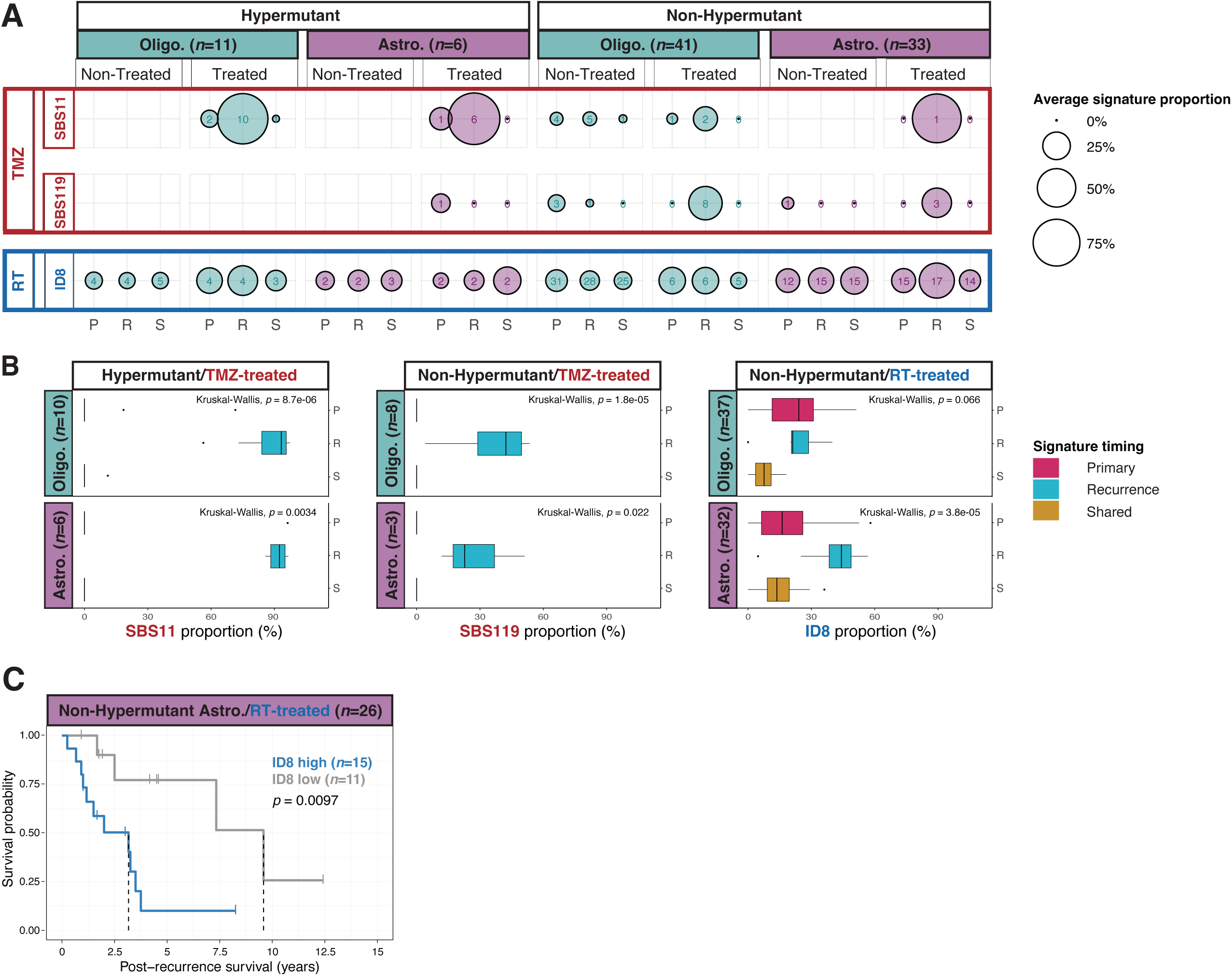
Mutational signatures SBS11 and SBS119 are associated with temozolomide exposure, while ID8 is linked to radiotherapy. SBS and ID signatures were compared using the average percentage of alterations attributed to these signatures (circle size) and the number of patients with any alteration attributable to these signatures (number in circle) per glioma type, hypermutation status, treatment regimen, and timing of the signatures. Treatment-associated signatures (**A:** landscape, **B:** highlights) were identified both glioma types. Prior treatment with temozolomide was associated with SBS11 in hypermutant gliomas and the newly identified SBS119 in non-hypermutant gliomas (oligodendrogliomas>astrocytomas). Recurrence-only fractions of radiotherapy-associated ID8 were increased in non-hypermutant astrocytomas only, though a similar (non-significant) trend was seen in non-hypermutant oligodendrogliomas. **C.** A high fraction of recurrence-only ID8 signature was correlated with poor prognosis. Abbreviations: Astro., astrocytomas; Oligo., oligodendrogliomas; TMZ, temozolomide; RT, radiotherapy; SBS, single-base substitutions; ID, small insertions and deletions; P, primary; R, recurrence; S, shared.

Radiation-treated astrocytomas that were not hypermutated showed significant enrichment of signature ID8 (**Fig. 6A and 6B**). While this signature was also detected in non-irradiated astrocytomas, it was highest in radiation-treated astrocytomas, suggesting that radiation increases the selection for ID8 mutations but not necessarily their formation. Radiation-treated oligodendrogliomas also evidenced an increase in ID8 mutations, but this trend did not reach statistical significance. The presence of the ID8 signature is significantly associated with inferior post-recurrence outcomes in both glioma subtypes (**Fig. 6C**). Altogether, the results from these mutational signatures further delineate the different evolutionary trajectories and responses to treatment between the two IDH-mutant glioma subtypes.

## Discussion

The IDH-mutant glioma subtypes oligodendroglioma and astrocytoma account for a disproportionate number of life years lost, emphasizing the need for novel therapeutic modalities^6^. A first step towards this goal was achieved with the recent FDA approval of IDH-inhibitor vorasidenib for the treatment of progressive WHO Grade II IDH-mutant gliomas in 2024. While this new treatment improves survival, the tumors eventually progress to higher grade and become resistant to the drug treatment^47^. The GLASS Consortium was established to characterize adaptive changes occurring in response to therapeutic treatments by comparing pre-and post-treatment gliomas with the goal of discovering new therapeutic vulnerabilities^13^. The current GLASS-IDH study offers the largest longitudinal analysis of IDH-mutant gliomas to date and for the first time, integrating whole-genome sequencing on a global, multi-institutional cohort spanning ten countries. The current dataset provides a substantial increase over previous releases^15,48^, both in IDH-mutant glioma patient numbers (from <100 to 206) and the addition of a large number of whole-genome datasets (*n* = 289 tumor samples) and the availability of detailed clinical annotation. The detailed clinical annotation of the GLASS cohort portrays the range of common treatment modalities over three decades with the preferential treatment comprising both radiotherapy and alkylating chemotherapy in the form of temozolomide for patients with both IDH-mutant tumor types.

Our analysis focused on subtype-stratified evolution, and we revealed evolutionary trajectories in both glioma subtypes by integrating clinical annotation and genomic findings in these significantly expanded datasets. This includes the detection of driver gene alterations that are enriched at recurrence (*PIK3CA*, *NOTCH1*, *MSH6* mutations in oligodendrogliomas; *CDKNA2* homozygous deletions, *PTPRZ1*/*MET*, *MYC/MYCN*, *CDK4*, *CCND2, EGFR, PDGFRA* focal amplifications in astrocytomas), which showed potential to help refine clinical classifications. This is of increased importance considering the limited vorasidenib indication. We further revealed non-specific gains in mutational and aneuploidy burden, and treatment-associated mutational signatures including SBS11, SBS119 and ID8. Considered together, a clear and dichotomous pattern emerged from these results. Oligodendrogliomas progress with new somatic single-nucleotide variants in driver genes as well as new single-base substitution signatures, while astrocytomas activate oncogenes and deactivate tumor suppressor genes by focal DNA copy number alterations which coincide with significant increases in aneuploidy burden (**Fig. 7**). Importantly, PI3K/*NOTCH1* mutations in non-hypermutant oligodendroglioma and acquired focal amplification of at least one well-described oncogene in non-hypermutated astrocytomas appeared to designate unfavorable outcome for, pointing to additional prognostic biomarkers to guide clinical management and extending previous observations^49^. Similarly, our findings provided further confirmation that alkylating agent-induced SBS11 mutational signature hypermutation is prognostically significant, with may impact current clinical practice by guiding follow-up therapy decisions (e.g., vorasidenib). Our analysis revealed that the IDH-mutant glioma subtypes further differed in treatment-related acquired mutational signatures, where recurrent astrocytomas acquired the radiation-associated ID8 signature while non-hypermutant recurrent oligodendrogliomas were enriched for the newly discovered temozolomide-induced SBS119 signature. Further studies, which may be supported by the availability of transcriptomic^48^ and methylomic^50^ datasets on subsets of the GLASS cohort, are needed to address new hypotheses arising from our results, such as whether the absence of the ID8 signature in oligodendroglioma suggests limited ability of oligodendrogliomas to derive radiotherapy resistance, why MSH6-mutant hypermutators show higher mutational burden, or if acquiring the SBS119 signature can be predicted.

**Figure 7.**
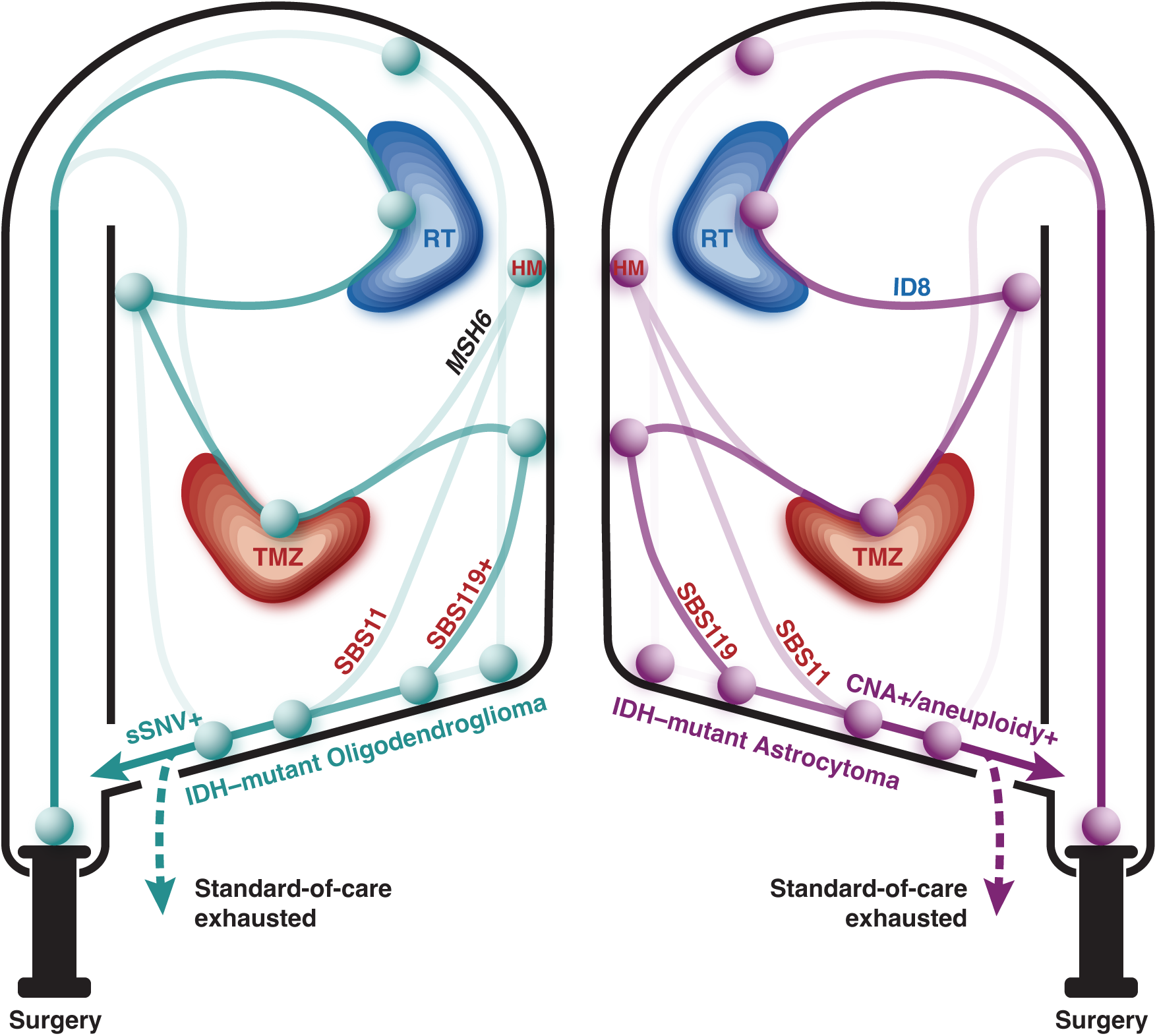
IDH-mutant gliomas evolve under shared pressures, yet the resulting alterations differ in type or extent. After surgery, most IDH-mutant gliomas are treated with radiotherapy and/or temozolomide. Though shaped by the same treatment-based selection pressures, changes in the genomes of astrocytomas and oligodendrogliomas suggest that different DNA damage response mechanisms are activated in response to therapy, resulting in acquiring of distinct sets of somatic alterations. At progression, astrocytomas are enriched for focal DNA copy number alterations, the ID8 signature and aneuploidies, whereas recurrent oligodendrogliomas evolve with new mutations in PI3K and *NOTCH1* genes, and higher rates of SBS119 after temozolomide exposure. The inevitable recurrence of disease either leads to a recurrence surgery and possible (re-)treatment with chemoradiation or a palliative setting when all standard-of-care options have been exhausted. Abbreviations: RT, radiotherapy; TMZ, temozolomide; HM, hypermutation; SBS, single-base substitutions; ID, small insertions and deletions; sSNV, somatic single nucleotide variant; CNA, copy number alteration.

The near-ubiquitous presence of *TP53* mutations in astrocytomas may contribute to progression mechanisms such as mitotic errors resulting in chromosomal missegregation events^51^, and ATRX deficiency has been repeatedly associated with replication stress, DNA damage, and chromosomal cohesion and congression abnormalities^52,53^. While the precise role of the 1p/19q in oligodendrogliomagenesis is not yet defined, the frequent presence of mutations in *CIC,* residing on 19q and which functions downstream of growth factor receptor signaling pathways, may provide one explanation for the absence of focal oncogene amplifications in oligodendrogliomas^54^. These observations highlight key differences in the response to treatment and drivers of tumor progression between the two subtypes of IDH-mutant glioma. Therapeutic strategies aimed at targeting genomic instability via KIF18A inhibition^36^ or related other vulnerabilities^55^ are likely to only be effective in astrocytomas. Future studies that provide new functional context to mutational signatures^56^ are needed to define the less-well understood mechanisms driving these genetic responses. The continued expansion of the GLASS datasets should further bolster understanding of optimally therapeutic design aimed at improving outcomes for patients with an IDH-mutant glioma.

In summary, astrocytomas and oligodendrogliomas undergo fundamentally distinct evolutionary trajectories in response to therapy, with astrocytomas accumulating copy number alterations, focal oncogene amplifications, and widespread aneuploidy, while oligodendrogliomas acquire new driver mutations and unique therapy-induced mutational signatures. These divergent patterns of genomic adaptation reflect subtype-specific DNA damage responses and provide a rationale for testing a precision oncology approach of targeting genomic instability or specific amplified oncogenes (e.g., *MET*) in astrocytomas, whereas oligodendrogliomas may be more responsive to pathway-directed therapies against acquired PI3K or *NOTCH1* mutations. This supports research to identify subtype-stratified and evolution-informed treatment strategies in IDH-mutant glioma. In addition, this work provides a vital framework required for future work looking at the molecular changes post vorasidenib treatment and resistance.

## Acknowledgements

We thank the patients and their families for the generous donations of their samples to biomedical research. This work is supported by NIH grants R01 CA237208, R01 CA271601, R21 CA256575, R21 NS114873 and Cancer Center Support Grant P30 CA034196; and a generous gift by the Dabbiere family (R.G.W.V); the Jane Coffin Childs Memorial Fund for Medical Research, JAX Scholar awards (F.S.V.). E.K. is supported by Emmy Noether Program (DFG, 541076190), Memorial Fellowship (Else Kröner-Fresenius-Stiftung). A.G. is supported by FNR INTER/TRANSCAN22/17612718/PLASTIG. A.I. is funded by R35CA253183. D.R.O. is supported by University of Colorado Department of Neurosurgery Nervous System Biorepository. I.D. and L.P. are supported by R01CA222146. J.M.C and P.S.L. are supported by R01CA218144 and Strain for the Brain 5K Run. J.S.B. is a full-time, paid employee of NIH/NCI. P.J.F. is supported by KWF (11026).

## Author contributions

TC, CMST, FV led data analysis and interpreted results in collaboration with EK, KCJ, JL, CH, CK, ZE, RGWV. Data processing is done by TC, CMST, FV with additional support from CH, GYY, TW, SBA, ES, KJA, HM. KCJ, EK, KJA, FV, PSG, ES, DMA, EBC, RGWV provided further insights into data analysis and interpretations. AL, AES, ARB, AG, AI, BH, CE, CH, CH, DRO, EGV, HKN, HM, JTH, JMC, JSB, JC, KM, KL, KW, LP, MPN, MP, MCMK, MS, MK, PW, PSL, PJF, RV, SN, SHP, WKAY, WRV, YH collected samples and/or clinical data. PLG helped with sample sequencing. TC, CMST, FV, RGWV led manuscript writing assisted by editing and revisions from EK, KCJ, AVC, BW, CW, DRO, DJB, EBC, EGV, GF, ID. JTH, LP. LG, MPN, MS, MW, MK, SP, TW, AES, ARB, AG, AMW, HKG, JMC, JSB, KA, LP, PSL, PJF, YH. TC, CMST, EK, KCJ, JL, ZE, CH, KJA, DMA, EBC, AI, DRO, DJB, EGV, GF, HKG, ID, KA, LG, MPN, MS, MK, PSL, PJF, SN, SHP, SP, FV, RGWV participated in internal discussions regarding data interpretation via offline meetings and/or teleconference. JL, ZE, CH, SBA, ES, KJA, AVC, AES, ARB, AG, AMW, AI, BW, CW, DJB, DMA, EBC, EGV, GF, HKG, ID, JTH, JMC, JSB, JC, KA, LP, LG, MPN, MCMK, MS, MW, MK, PSL, PJF, SN, SHP, SP, TW, WKAY, FV, RGWV are GLASS consortium members. RGWV led the GLASS consortium and this project.

## Methods

### DNA sequencing

Whole-genome sequencing (WGS) and whole-exome sequencing (WXS) data were newly generated for MD Anderson Cancer Center, Duke University Hospital, Henry Ford Hospital, Case Western Reserve University, Chinese University of Hong Kong, Mayo Clinic, Northwestern Medicine, Olivia Newton-John Cancer Research Institute, University of California San Francisco, Saint Joseph’s Hospital and Medical Center, Seoul National University, and Luxembourg Institute of Health at the Jackson Laboratory for Genome Medicine sequencing core. All were subjected to 150 bases paired-end sequencing using Novaseq (Illumina, CA, USA), according to manufacturer’s protocol. Previously published longitudinal data from GLASS^15,48^ and the Cancer Genome Atlas (TCGA)^4^ were also used and the sequencing information for those can be found at previous publications. In total, sequencing data was contributed by 21 institutions: Columbia University, University of Florida, German Cancer Research Center, Pitié-Salpêtrière, Samsung Medical Center, Biobank Japan, University of South Wales, University of Sao Paulo, and the aforementioned 13 institutions including the Jackson Laboratory. For the Seoul National University cohort, both DNA and RNA were concurrently extracted from each tumor sample at The Jackson Laboratory for Genomic Medicine using Qiagen’s AllPrep DNA/RNA Mini Kit (#80204). For blood samples, DNA extraction was carried out using Qiagen’s QIAamp DNA Mini and Blood Mini Kit (#51104). A total of 200 nanograms of DNA was fragmented to 400 base pairs using a LE220 focused-ultrasonicator (Covaris) and then size-selected with Ampure XP beads (Beckman Coulter). The DNA fragments underwent end-repair, A-tailing, and the ligation of Illumina unique adapters (Illumina) using the KAPA Hyper Prep Kit from Roche (#7962363001). The resulting whole genome libraries were sequenced with 150 base pair paired-end reads on the Illumina NovaSeq 6000, achieving 25X coverage for normal samples and greater than 35-40X for tumor samples. RNA sequencing libraries were prepared using the KAPA mRNA Hyperprep Kit from Roche (#8098123702) and sequenced on the Illumina NovaSeq 6000 platform, producing 150 base pair paired-end reads.

### Reference cohorts

GLASS data was compared to two reference cohorts: TCGA^4^ and the American Association for Cancer Research (AACR) Project Genomics Evidence Neoplasia Information Exchange (GENIE), v18.5^57^. Clinical data of GLASS patients at baseline was compared to TCGA patients with primary IDH-mutant gliomas with respect to age of onset and overall survival (OS)^4,15,48^. Survival data of patients with oligodendrogliomas and astrocytomas were compared between the GLASS and GENIE cohorts^57^. The GENIE cohort includes mainly panel sequencing data, and in most cases lacks copy number data or coverage of *TERT* promoter hotspots. Oligodendrogliomas in the GENIE cohort were therefore defined as IDH-mutant gliomas without alterations in *TP53* or *ATRX* as determined by high-coverage panel sequencing, whereas IDH-mutant gliomas with these alterations were called astrocytomas.

### GLASS identifiers

GLASS barcode system based on the TCGA barcode design were used to identify all samples and their corresponding patients. Briefly the 24-character barcode includes information for the project, collection center, patient identification, sample type, analyte type and sequencing platform. The detailed information for the barcode breakdown is available in the previous GLASS publication^48^.

### Computational pipelines

GLASS pipelines for variant and copy number calling were developed using Snakemake (v5.2.2)^58^. Genome Analysis Toolkit 4 (GATK4, v4.1.0.0)^59^ suite was used unless otherwise specified. All data were analyzed using homogenous pipelines capable of processing raw FastQ files.

### Alignment and variant calling

Briefly, in alignment and preprocessing, the study adhered to GATK Best Practices, using various tools to manage and sanitize Binary Alignment Map (BAM) files, assign, and manage read groups, perform quality control, mark sequencing adapters, align sequences to the b37/hg19 genome (human_g1k_v37_decoy), merge and mark duplicates, and perform base recalibration. For variant detection, GATK4 was used, with germline variants called from control samples, and somatic variants identified in tumor samples and patient cohorts. Quality control measures included evaluating cross-sample contamination and read orientation artifacts. In variant post-processing, BCFTools (v1.9)^60^ normalized, sorted, and indexed variants to generate a consensus Variant Call Format (VCF) file, which was then annotated and used to compare allele frequencies between related samples, with single-sample variants overlaid on multi-sample calls to infer individual sample mutations. Further details of these steps can be found in the previously published GLASS paper^15^.

### Copy number calling

Copy number identification was performed according to GATK Best Practices, with different methods for WGS and WXS data. For WGS data, the genome was segmented into 10kb bins, while overlapping regions from various capture kits were merged for WXS data. Both strategies included counting reads in these bins, subsetting germline variants, and separating the cohort by sequencing center to create a panel of normals. This involved denoising read counts, annotating intervals for GC content, and inspecting results for quality. Copy number calling utilized a method based on GATK ‘CallCopyRatioSegments’ and GISTIC (v2.0.23)^61,62^, identifying neutral segments and classifying amplifications or deletions based on weighted means and standard deviations. Chromosome arms were analyzed to set high-level thresholds, and gene-level copy numbers were called by intersecting gene boundaries with segment intervals and comparing the weighted non-log2 copy ratio to these thresholds. These steps are also described in detail previously^15^.

### dNdScv

The dN/dS ratios were estimated using dNdScv (v0.1.0)^63^ with the default and recommended parameters applied to all mutations in initial and recurrent tumor samples, as well as for each mutational fraction (unique to initial, unique to recurrent, and shared). These analyses were performed separately within each of the two IDH-mutant glioma types.

### MutSigCV

To identify significantly mutated genes during tumor evolution, we used MutSigCV (v1.3.5)^37^ on WXS and WGS data from IDH-mutant astrocytomas and oligodendrogliomas. Separate analyses were conducted for primary and recurrent samples to detect acquired driver mutations. Recurrent tumors were further classified into hypermutated and non-hypermutated groups, with MutSigCV applied independently to each group. Genes lacking sufficient coverage or associated with known germline variants were excluded, and statistical significance was determined using MutSigCV-calculated false discovery rates (FDR).

### GISTIC

GISTIC^61^ was run on the public version of the GenePattern server^64^, on the segmentation data of primary and recurrent tumors for each IDH-mutant glioma type individually to identify broad and focal copy number changes of significance in each group. The analysis was conducted using the GeneGISTIC method with broad event detection and gene collapse values set to extreme. Additional parameters included a focal length cutoff ≥ 0.5 with a confidence level of 99% and a chromosome arm adjustment capped to 1.0. Copy number driver genes were identified from the GeneGISTIC results.

### Aneuploidy calculation

Aneuploidy was calculated at three different levels for all samples, arm-level, segment level, and genome-wide level.

For the arm-level aneuploidy calculation, a previously described method was used^65^. Briefly, adjacent segments with identical arm-level calls (−1, 0, or 1) were merged, resulting in a single call for each merged segment. The proportion of the chromosome arm spanned by each merged segment was calculated. Segments covering over 80% of the arm were called −1 (loss), 0 (neutral), +1 (gain), or ‘NA’ if no segment covered at least 80%. For each sample, the number of arms with non-neutral events was then counted. As a secondary method, average log2(copy number ratios) were calculated to compare arm-level aneuploidies between groups. Lower thresholds were set at ±0.1 for significant arm-level gain or loss at any timepoint and ±0.05 for significant arm-level changes (Δ gains or losses).

For segment level calculation, each segment was assigned a status of gain, loss, or neutral based on the log_₂_ copy ratio threshold of ±0.2. Segments with absolute log_₂_ ratio > 0.2 were considered altered. each sample, the total number of altered segments (gains and losses) was counted.

For genome-wide level calculation, fraction of genome altered was calculated with the following formula: fraction of genome altered = 1 – (total base pairs in neutral segments/total callable base pairs). For WGS samples, the callable genome size was assumed to be 3×10D bp. For WXS samples, callable size was set to 5×10D bp based on the exome capture target region.

### Mutational signatures

Mutational signature analysis was done in three steps using previously published methods^39,66,67^. First, WGS data for all the samples were used to create a mutational matrix and extract *de novo* signatures using SigProfilerExtractor tool (v1.2.1)^66^. We used the b37/hg19 reference genome, maximum signatures of 10 and non-negative matrix factorization (NMF) replicates of 100 for the SigProfiler run. Then, *de novo* signatures were deconvoluted to a curated list of reference signatures for single-base substitutions (SBS), double-base substitutions (DBS) and small insertions and deletions (ID). To curate the reference signature list, we combined two public mutational signature databases listed; namely the Catalogue Of Somatic Mutations In Cancer (COSMIC) database (cancer.sanger.ac.uk/signatures)^39^ and the Signal database (signal.mutationalsignatures.com/explore/main/cancer/signatures)^40^ signatures. To ensure confidence in signature assignment probability, we used only WGS data from our cohort, aligning with the WGS-based curated reference. Finally, we used the deconvoluted reference signatures identified in step 2 to estimate the exposure of mutational signatures in each sample. The R package Palimpsest (v2.0.0)^68^ was then employed to assign the most likely signature to each mutation within a sample.

### Mutational burden

Mutational burden was determined by the number of mutations per Mb sequenced, requiring a minimum base coverage of 15X. DNA hypermutation was defined for recurrent tumors with more than 10 mutations per Mb and having evidence of COSMIC mutational signature SBS11.

### Mutual exclusivity analysis

Mutual exclusivity of genomic alterations was assessed using the Discrete Independence Statistic Controlling for Observations with Varying Event Rates (DISCOVER) algorithm (v3.1.7)^23^, which computes tumor-specific probabilities of gene alterations while preserving the observed marginal totals per gene and per tumor. Expected co-occurrence rates between gene pairs were derived under a Poisson–binomial model, accounting for these probabilities. Significantly fewer co-alterations than expected under this null model were interpreted as evidence of mutual exclusivity.

### Survival analysis

Survival endpoints included OS from diagnosis and time to recurrence, measured from the date of initial surgical resection to the date of event (death or recurrence/progression) or censored at last follow-up. OS from birth was calculated by summing age at diagnosis and OS from diagnosis. Post-recurrence survival was defined from the date of recurrence surgery to death or censoring. Kaplan–Meier survival curves were generated and compared using the log-rank test with the survival and survminer R packages. Cox proportional hazards (PH) regression was used to estimate hazard ratios (HRs); variables with likelihood ratio test *p* values ≤ 0.10 were considered for inclusion in multivariable models. Schoenfeld residuals were examined to verify the PH assumption; variables violating this assumption were excluded. Prognostic models were refined through stepwise backward elimination, and model performance was quantified using Harrell’s concordance index (C-index) with 1000 bootstrap resamples.

### Other Statistical Methods

All statistical analyses were performed using R (v4.3.2) and Python (v3.10.13). Unpaired categorical variables were compared using Fisher’s exact test, and unpaired continuous variables with the Wilcoxon rank-sum test or the Kruskal–Wallis test, as appropriate. Paired categorical variables were compared using McNemar’s test. Statistical significance was defined as a two-tailed *p* value < 0.05 unless specified otherwise.

## Data Availability

Deidentified somatic variant and clinical datasets are publicly available via Synapse (http://synapse.org) with Project SynID syn17038081.

## Code Availability

Custom computational pipeline scripts for sequencing data pre-processing are accessible via the project’s GitHub repository (https://github.com/fpbarthel/GLASS). Downstream data analysis scripts for this study will be made available on github at the time of publication.

## References

1. Price, M., et al. CBTRUS Statistical Report: Primary Brain and Other Central Nervous System Tumors Diagnosed in the United States in 2017-2021. Neuro Oncol 26, vi1-vi85 (2024). 10.1093/neuonc/noae145

2 Louis, D. N. et al. The 2021 WHO Classification of Tumors of the Central Nervous System: a summary. Neuro Oncol 23, 1231–1251 (2021). 10.1093/neuonc/noab106

3 Miller, J. J. et al. Isocitrate dehydrogenase (IDH) mutant gliomas: A Society for Neuro-Oncology (SNO) consensus review on diagnosis, management, and future directions. Neuro Oncol 25, 4–25 (2023). 10.1093/neuonc/noac207

4. Cancer Genome Atlas Research, N., et al. Comprehensive, Integrative Genomic Analysis of Diffuse Lower-Grade Gliomas. N Engl J Med 372, 2481-2498 (2015). 10.1056/NEJMoa1402121

5 Eckel-Passow, J. E. et al. Glioma Groups Based on 1p/19q, IDH, and TERT Promoter Mutations in Tumors. N Engl J Med 372, 2499–2508 (2015). 10.1056/NEJMoa1407279

6 Gerstl, J. V. E. et al. Years of life lost due to central nervous system tumor subtypes in the United States. Neuro Oncol (2025). 10.1093/neuonc/noaf142

7 Touat, M. et al. Mechanisms and therapeutic implications of hypermutation in gliomas. Nature 580, 517–523 (2020). 10.1038/s41586-020-2209-9

8 Kocakavuk, E. et al. Radiotherapy is associated with a deletion signature that contributes to poor outcomes in patients with cancer. Nat Genet 53, 1088–1096 (2021). 10.1038/s41588-021-00874-3

9 Claus, E. B. & Verhaak, R. G. W. Targeting IDH in Low-Grade Glioma. N Engl J Med 389, 655–659 (2023). 10.1056/NEJMe2305602

10 Mellinghoff, I. K. et al. Vorasidenib in IDH1-or IDH2-Mutant Low-Grade Glioma. N Engl J Med 389, 589–601 (2023). 10.1056/NEJMoa2304194

11 Parsons, D. W. et al. An integrated genomic analysis of human glioblastoma multiforme. Science 321, 1807–1812 (2008). 10.1126/science.1164382

12 Wang, F. et al. Leukemia stemness and co-occurring mutations drive resistance to IDH inhibitors in acute myeloid leukemia. Nat Commun 12, 2607 (2021). 10.1038/s41467-021-22874-x

13 Consortium, G. Glioma through the looking GLASS: molecular evolution of diffuse gliomas and the Glioma Longitudinal Analysis Consortium. Neuro Oncol 20, 873–884 (2018). 10.1093/neuonc/noy020

14 Aldape, K. et al. Glioma Through the Looking GLASS: Molecular Evolution of Diffuse Gliomas and the Glioma Longitudinal AnalySiS Consortium. Neuro-Oncology, noy020-noy020 (2018). 10.1093/neuonc/noy020

15 Barthel, F. P. et al. Longitudinal molecular trajectories of diffuse glioma in adults. Nature 576, 112–120 (2019). 10.1038/s41586-019-1775-1

16 Johnson, B. E. et al. Mutational analysis reveals the origin and therapy-driven evolution of recurrent glioma. Science 343, 189–193 (2014). 10.1126/science.1239947

17 Mu, Q. et al. Identifying predictors of glioma evolution from longitudinal sequencing. Sci Transl Med 15, eadh4181 (2023). 10.1126/scitranslmed.adh4181

18 Kocakavuk, E. et al. Hemizygous CDKN2A deletion confers worse survival outcomes in IDHmut-noncodel gliomas. Neuro Oncol 25, 1721–1723 (2023). 10.1093/neuonc/noad095

19 Shi, D. D., Anand, S., Abdullah, K. G. & McBrayer, S. K. DNA damage in IDH-mutant gliomas: mechanisms and clinical implications. J Neurooncol 162, 515–523 (2023). 10.1007/s11060-022-04172-8

20 Ceccarelli, M. et al. Molecular Profiling Reveals Biologically Discrete Subsets and Pathways of Progression in Diffuse Glioma. Cell 164, 550–563 (2016). 10.1016/j.cell.2015.12.028

21 van den Bent, M. J. & Chang, S. M. Grade II and III Oligodendroglioma and Astrocytoma. Neurol Clin 36, 467–484 (2018). 10.1016/j.ncl.2018.04.005

22 Johnson, B. E. et al. Mutational analysis reveals the origin and therapy-driven evolution of recurrent glioma. Science 343, 189–193 (2014). 10.1126/science.1239947

23 Canisius, S., Martens, J. W. & Wessels, L. F. A novel independence test for somatic alterations in cancer shows that biology drives mutual exclusivity but chance explains most co-occurrence. Genome Biol 17, 261 (2016). 10.1186/s13059-016-1114-x

24 Lee, Y. R., Chen, M. & Pandolfi, P. P. The functions and regulation of the PTEN tumour suppressor: new modes and prospects. Nat Rev Mol Cell Biol 19, 547–562 (2018). 10.1038/s41580-018-0015-0

25 Palomero, T. et al. Mutational loss of PTEN induces resistance to NOTCH1 inhibition in T-cell leukemia. Nat Med 13, 1203–1210 (2007). 10.1038/nm1636

26 Niu, N. et al. Tumor cell-intrinsic epigenetic dysregulation shapes cancer-associated fibroblasts heterogeneity to metabolically support pancreatic cancer. Cancer cell 42, 869–884 e869 (2024). 10.1016/j.ccell.2024.03.005

27 Halani, S. H. et al. Multi-faceted computational assessment of risk and progression in oligodendroglioma implicates NOTCH and PI3K pathways. NPJ Precis Oncol 2, 24 (2018). 10.1038/s41698-018-0067-9

28 Consortium, A. P. G. AACR Project GENIE: Powering Precision Medicine through an International Consortium. Cancer Discov 7, 818–831 (2017). 10.1158/2159-8290.CD-17-0151

29 Bao, Z. et al. PTPRZ1-METFUsion GENe (ZM-FUGEN) trial: study protocol for a multicentric, randomized, open-label phase II/III trial. Chin Neurosurg J 9, 21 (2023). 10.1186/s41016-023-00329-0

30 Bao, Z. S. et al. RNA-seq of 272 gliomas revealed a novel, recurrent PTPRZ1-MET fusion transcript in secondary glioblastomas. Genome Res 24, 1765–1773 (2014). 10.1101/gr.165126.113

31 Hu, H. et al. Mutational Landscape of Secondary Glioblastoma Guides MET-Targeted Trial in Brain Tumor. Cell 175, 1665–1678 e1618 (2018). 10.1016/j.cell.2018.09.038

32 Yu, Y. et al. Temozolomide-induced hypermutation is associated with distant recurrence and reduced survival after high-grade transformation of low-grade IDH-mutant gliomas. Neuro Oncol 23, 1872–1884 (2021). 10.1093/neuonc/noab081

33 Galbraith, K. et al. Prognostic value of DNA methylation subclassification, aneuploidy, and CDKN2A/B homozygous deletion in predicting clinical outcome of IDH mutant astrocytomas. Neuro Oncol 26, 1042–1051 (2024). 10.1093/neuonc/noae009

34 Mirchia, K. et al. Total copy number variation as a prognostic factor in adult astrocytoma subtypes. Acta Neuropathol Commun 7, 92 (2019). 10.1186/s40478-019-0746-y

35 Davoli, T., Uno, H., Wooten, E. C. & Elledge, S. J. Tumor aneuploidy correlates with markers of immune evasion and with reduced response to immunotherapy. Science 355 (2017). 10.1126/science.aaf8399

36 Zerbib, J., Bloomberg, A. & Ben-David, U. Targeting vulnerabilities of aneuploid cells for cancer therapy. Trends Cancer (2025). 10.1016/j.trecan.2025.04.005

37 Lawrence, M. S. et al. Mutational heterogeneity in cancer and the search for new cancer-associated genes. Nature 499, 214–218 (2013). 10.1038/nature12213

38 Braganza, M. Z. et al. Ionizing radiation and the risk of brain and central nervous system tumors: a systematic review. Neuro Oncol 14, 1316–1324 (2012). 10.1093/neuonc/nos208

39 Alexandrov, L. B. et al. The repertoire of mutational signatures in human cancer. Nature 578, 94–101 (2020). 10.1038/s41586-020-1943-3

40 Degasperi, A. et al. Substitution mutational signatures in whole-genome-sequenced cancers in the UK population. Science 376 (2022). 10.1126/science.abl9283

41 Singh, V. K., Rastogi, A., Hu, X., Wang, Y. & De, S. Mutational signature SBS8 predominantly arises due to late replication errors in cancer. Commun Biol 3, 421 (2020). 10.1038/s42003-020-01119-5

42 Jin, H. et al. Accurate and sensitive mutational signature analysis with MuSiCal. Nat Genet 56, 541–552 (2024). 10.1038/s41588-024-01659-0

43 Fukagawa, A. et al. Genomic and epigenomic integrative subtypes of renal cell carcinoma in a Japanese cohort. Nat Commun 14, 8383 (2023). 10.1038/s41467-023-44159-1

44 Senkin, S. et al. Geographic variation of mutagenic exposures in kidney cancer genomes. Nature 629, 910–918 (2024). 10.1038/s41586-024-07368-2

45 Thatikonda, V. et al. Comprehensive analysis of mutational signatures reveals distinct patterns and molecular processes across 27 pediatric cancers. Nat Cancer 4, 276–289 (2023). 10.1038/s43018-022-00509-4

46 Liu, K. & Jiang, Y. Polymorphisms in DNA Repair Gene and Susceptibility to Glioma: A Systematic Review and Meta-Analysis Based on 33 Studies with 15 SNPs in 9 Genes. Cell Mol Neurobiol 37, 263–274 (2017). 10.1007/s10571-016-0367-y

47 Darlix, A. et al. Who will benefit from vorasidenib? Review of data from the literature and open questions. Neurooncol Pract 12, i6–i18 (2025). 10.1093/nop/npae104

48 Varn, F. S. et al. Glioma progression is shaped by genetic evolution and microenvironment interactions. Cell 185, 2184–2199 e2116 (2022). 10.1016/j.cell.2022.04.038

49 Yang, R. R. et al. IDH mutant lower grade (WHO Grades II/III) astrocytomas can be stratified for risk by CDKN2A, CDK4 and PDGFRA copy number alterations. Brain Pathol 30, 541–553 (2020). 10.1111/bpa.12801

50 Malta, T. M. et al. The Epigenetic Evolution of Glioma Is Determined by the IDH1 Mutation Status and Treatment Regimen. Cancer Res 84, 741–756 (2024). 10.1158/0008-5472.CAN-23-2093

51 Zhao, M. et al. Mutant p53 gains oncogenic functions through a chromosomal instability-induced cytosolic DNA response. Nat Commun 15, 180 (2024). 10.1038/s41467-023-44239-2

52 Lovejoy, C. A. et al. Loss of ATRX, genome instability, and an altered DNA damage response are hallmarks of the alternative lengthening of telomeres pathway. PLoS Genet 8, e1002772 (2012). 10.1371/journal.pgen.1002772

53 Wang, Y. et al. G-quadruplex DNA drives genomic instability and represents a targetable molecular abnormality in ATRX-deficient malignant glioma. Nat Commun 10, 943 (2019). 10.1038/s41467-019-08905-8

54 Cahill, D. P., Louis, D. N. & Cairncross, J. G. Molecular background of oligodendroglioma: 1p/19q, IDH, TERT, CIC and FUBP1. CNS Oncol 4, 287–294 (2015). 10.2217/cns.15.32

55 Girish, V. et al. Oncogene-like addiction to aneuploidy in human cancers. Science 381, eadg4521 (2023). 10.1126/science.adg4521

56 Kucab, J. E. et al. A Compendium of Mutational Signatures of Environmental Agents. Cell 177, 821–836 e816 (2019). 10.1016/j.cell.2019.03.001

57 Pugh, T. J. et al. AACR Project GENIE: 100,000 Cases and Beyond. Cancer Discov 12, 2044–2057 (2022). 10.1158/2159-8290.CD-21-1547

58 Koster, J. & Rahmann, S. Snakemake-a scalable bioinformatics workflow engine. Bioinformatics 34, 3600 (2018). 10.1093/bioinformatics/bty350

59 Van der Auwera, G. A. et al. From FastQ data to high confidence variant calls: the Genome Analysis Toolkit best practices pipeline. Curr Protoc Bioinformatics 43, 11 10 11–11 10 33 (2013). 10.1002/0471250953.bi1110s43

60 Li, H. et al. The Sequence Alignment/Map format and SAMtools. Bioinformatics 25, 2078–2079 (2009). 10.1093/bioinformatics/btp352

61 Mermel, C. H. et al. GISTIC2.0 facilitates sensitive and confident localization of the targets of focal somatic copy-number alteration in human cancers. Genome Biol 12, R41 (2011). 10.1186/gb-2011-12-4-r41

62 Beroukhim, R. et al. Assessing the significance of chromosomal aberrations in cancer: methodology and application to glioma. Proc Natl Acad Sci U S A 104, 20007–20012 (2007). 10.1073/pnas.0710052104

63 Martincorena, I. et al. Universal Patterns of Selection in Cancer and Somatic Tissues. Cell 171, 1029–1041 e1021 (2017). 10.1016/j.cell.2017.09.042

64 Reich, M. et al. GenePattern 2.0. Nat Genet 38, 500–501 (2006). 10.1038/ng0506-500

65 Taylor, A. M. et al. Genomic and Functional Approaches to Understanding Cancer Aneuploidy. Cancer Cell 33, 676–689 e673 (2018). 10.1016/j.ccell.2018.03.007

66 Islam, S. M. A. et al. Uncovering novel mutational signatures by de novo extraction with SigProfilerExtractor. Cell Genom 2, None (2022). 10.1016/j.xgen.2022.100179

67 Letouze, E. et al. Mutational signatures reveal the dynamic interplay of risk factors and cellular processes during liver tumorigenesis. Nat Commun 8, 1315 (2017). 10.1038/s41467-017-01358-x

68 Shinde, J. et al. Palimpsest: an R package for studying mutational and structural variant signatures along clonal evolution in cancer. Bioinformatics 34, 3380–3381 (2018). 10.1093/bioinformatics/bty388

