## Supplementary Tables for "Tumor-initiating genetics and therapy drive divergent molecular evolution in IDH-mutant gliomas"

| | IDH-mutant Astrocytoma | IDH-mutant Oligodendroglioma |
|---|---|---|
| **Clinical markers** | | |
| *Demographics* | | |
| Age of diagnosis (>40 vs ≤40) | 1.64 [1.00-2.71] | **2.31 [1.25-4.27]** |
| Sex (male vs female) | 1.30 [0.83-2.03] | 1.39 [0.74-2.61] |
| *Primary treatment* | | |
| Surgery type (resection vs biopsy) | 1.79 [0.54-5.96] | 1.29 [0.46-3.61] |
| Radiotherapy (yes vs no) | **2.14 [1.36-3.36]** | **5.13 [2.44-10.81]** |
| Alkylating chemotherapy (yes vs no) | 1.36 [0.86-2.15] | **2.67 [1.34-5.33]** |
| *Primary pathology* | | |
| Grade (3 vs other)** | 1.67 [0.90-3.10] | 9.20 [2.98-28.37]* |
| Grade (4 vs other)** | **3.03 [1.61-5.68]** | |
| **Molecular markers at primary surgery** | | |
| *Mutations (present vs absent)* | | |
| *MSH6* | *all wildtype* | *all wildtype* |
| *NOTCH1* | **12.85 [2.17-75.94]** | 3.09 [0.87-10.94] |
| *PIK3CA* | **15.53 [2.56-94.10]** | 1.94 [0.67-5.66] |
| *PIK3R1* | *all wildtype* | 0.58 [0.09-3.68] |
| *Homozygous deletions (HD vs non-HD)* | | |
| *CDKN2A/B* | **2.28 [1.04-4.99]** | *all non-HD* |
| *PTEN* | 1.55 [0.29-8.23] | *all non-HD* |
| *ATRX* | 0.43 [0.13-1.43] | *all non-HD* |
| *Amplifications (amp vs non-amp)* | | |
| *CCND2* | **24.61 [7.65-79.16]** | *all non-amp* |
| *EGFR* | 2.30 [0.85-6.21] | <0.001 [0-∞] |
| *PDGFRA* | **100.70 [13.10-773.70]** | *all non-amp* |
| *CDK4/6* | **4.86 [2.20-10.74]** | *all non-amp* |
| *MET/PTPRZ1* | 0.99 [0.30-3.27] | 1.71 [0.31-9.41] |
| *MYC/MYCN* | **3.13 [1.42-6.89]** | *all non-amp* |
| *Methylation (methylated vs unmethylated)* | | |
| *MGMT* promoter | 0.68 [0.29-1.58]* | *all methylated* |

**Extended Data Table 1. Univariable Cox proportional hazards models concerning overall survival of GLASS IDH-mutant glioma patients.**
Hazard ratios with 90% confidence intervals. Significant factors for multivariable testing were defined as all variables with $p < 0.10$ that did not violate the proportional hazards assumption. *Proportional hazards assumption violated: Schoenfeld test $p < 0.05$. **Variable consist of 3 separate levels. Abbreviations: HD, homozygous deletion; amp, amplification.

| | IDH-mutant Astrocytoma | IDH-mutant Oligodendroglioma |
|---|---|---|
| **Full Multivariable Cox PH Model** | C-index = 0.69 | C-index = 0.79 |
| Age of diagnosis (>40 vs ≤40) | | **3.94 [1.58-9.80]** |
| Radiotherapy (yes vs no) | **2.25 [1.18-4.29]** | **5.54 [2.21-13.88]** |
| Alkylating chemotherapy (yes vs no) | | 2.02 [0.83-4.88] |
| Grade (3 vs other)** | 0.67 [0.27-1.65] | |
| Grade (4 vs other)** | 1.37 [0.45-4.17] | |
| *NOTCH1* (mutant vs wildtype) | **56.64 [5.41-593.05]** | |
| *PIK3CA* (mutant vs wildtype) | **75.78 [6.77-847.97]** | |
| *CDKN2A/B* (HD vs non-HD) | 1.59 [0.49-5.14] | |
| *CCND2* (amp vs non-amp) | 6.02 [0.55-65.55] | |
| *PDGFRA* (amp vs non-amp) | 16.77 [0.28-1002.35]* | |
| *CDK4/6* (amp vs non-amp) | 2.79 [0.73-10.71] | |
| *MYC/MYCN* (amp vs non-amp) | 1.12 [0.27-4.64] | |
| **Stepwise Backward Cox PH Model** | C-index = 0.66 | C-index = 0.79 |
| Age of diagnosis (>40 vs ≤40) | | **3.94 [1.58-9.80]** |
| Radiotherapy (yes vs no) | **1.99 [1.10-3.62]** | **5.54 [2.21-13.88]** |
| Alkylating chemotherapy (yes vs no) | | 2.02 [0.83-4.88] |
| *NOTCH1* (mutant vs wildtype) | **43.73 [4.46-429.04]** | |
| *PIK3CA* (mutant vs wildtype) | **55.26 [5.42-563.56]** | |
| *CCND2* (amp vs non-amp) | **23.89 [5.19-109.90]** | |
| *CDK4/6* (amp vs non-amp) | **3.25 [1.19-8.91]** | |

**Extended Data Table 2. Multivariable Cox proportional hazards models concerning overall survival of GLASS IDH-mutant glioma patients.**
Hazard ratios with 95% confidence intervals. Significant factors were defined as all variables with $p < 0.05$ that did not violate the proportional hazards assumption. Model performance was quantified using Harrell's concordance index (C-index). *Proportional hazards assumption violated: Schoenfeld test $p < 0.05$. **Variable consist of 3 separate levels. Abbreviations: PH, proportional hazards; HD, homozygous deletion; amp, amplification.

| | IDH-mutant Astrocytoma | IDH-mutant Oligodendroglioma |
|---|---|---|
| **Clinical markers** | | |
| *Demographics* | | |
| Age of diagnosis (>40 vs ≤40) | **1.67 [1.12-2.50]** | 1.01 [0.72-1.41] |
| Sex (male vs female) | 1.05 [0.74-1.49] | **1.99 [1.37-2.89]** |
| *Primary treatment* | | |
| Surgery type (resection vs biopsy) | 1.12 [0.51-2.43] | 0.71 [0.41-1.25] |
| Radiotherapy (yes vs no) | 0.86 [0.61-1.22] | 0.75 [0.49-1.13] |
| Alkylating chemotherapy (yes vs no) | **0.68 [0.47-0.98]** | 1.00 [0.67-1.49] |
| *Primary pathology* | | |
| Grade (3 vs other)** | 0.86 [0.51-1.44] | **2.25 [1.10-4.59]** |
| Grade (4 vs other)** | 1.23 [0.72-2.12] | |
| **Molecular markers at primary surgery** | | |
| *Mutations (present vs absent)* | | |
| *MSH6* | *all wildtype* | *all wildtype* |
| *NOTCH1* | 3.86 [0.71-20.89] | 1.20 [0.51-2.83] |
| *PIK3CA* | **9.37 [1.64-53.61]** | 1.76 [0.98-3.15] |
| *PIK3R1* | *all wildtype* | 0.82 [0.25-2.69] |
| *Homozygous deletions (HD vs non-HD)* | | |
| *CDKN2A/B* | 1.30 [0.64-2.63] | *all non-HD* |
| *PTEN* | 0.70 [0.13-3.68] | *all non-HD* |
| *ATRX* | 0.65 [0.28-1.51] | *all non-HD* |
| *Amplifications (amp vs non-amp)* | | |
| *CCND2* | **5.30 [1.91-14.67]** | *all non-amp* |
| *EGFR* | 2.25 [0.95-5.30] | 0.28 [0.05-1.50] |
| *PDGFRA* | **8.84 [2.50-31.25]** | *all non-amp* |
| *CDK4/6* | **2.25 [1.04-4.87]** | *all non-amp* |
| *MET/PTPRZ1* | 0.83 [0.31-2.20]* | 0.71 [0.26-1.92] |
| *MYC/MYCN* | 1.53 [0.75-3.09] | *all non-amp* |
| *Methylation (methylated vs unmethylated)* | | |
| *MGMT* promoter | 1.22 [0.58-2.55] | *all methylated* |

**Extended Data Table 3. Univariable Cox proportional hazards models concerning time to recurrence of GLASS IDH-mutant glioma patients.**
Hazard ratios with 90% confidence intervals. Significant factors for multivariable testing were defined as all variables with $p < 0.10$ that did not violate the proportional hazards assumption. *Proportional hazards assumption violated: Schoenfeld test $p < 0.05$. **Variable consist of 3 separate levels. Abbreviations: HD, homozygous deletion; amp, amplification.

| | IDH-mutant Astrocytoma | IDH-mutant Oligodendroglioma |
|---|---|---|
| **Full Multivariable Cox PH Model** | C-index = 0.62 | C-index = 0.58 |
| Age of diagnosis (>40 vs ≤40) | 1.61 [0.83-3.15] | |
| Sex (male vs female) | | **1.87 [1.12-3.13]** |
| Alkylating chemotherapy (yes vs no) | 0.63 [0.38-1.02] | |
| Grade (3 vs other) | | 1.68 [0.70-4.04] |
| *PIK3CA* (mutant vs wildtype) | **7.13 [0.79-64.43]** | |
| *CCND2* (amp vs non-amp) | 2.91 [0.38-22.08]* | |
| *PDGFRA* (amp vs non-amp) | 2.03 [0.11-36.13] | |
| *CDK4/6* (amp vs non-amp) | 2.29 [0.51-10.31] | |
| **Stepwise Backward Cox PH Model** | C-index = 0.61 | C-index = 0.57 |
| Age of diagnosis (>40 vs ≤40) | 1.82 [0.97-3.39] | |
| Sex (male vs female) | | **1.99 [1.28-3.11]** |
| Alkylating chemotherapy (yes vs no) | 0.62 [0.39-1.00] | |
| *PDGFRA* (amp vs non-amp) | **11.76 [2.50-55.35]** | |

**Extended Data Table 4. Multivariable Cox proportional hazards models concerning time to recurrence of GLASS IDH-mutant glioma patients.**
Hazard ratios with 95% confidence intervals. Significant factors were defined as all variables with $p < 0.05$ that did not violate the proportional hazards assumption. Model performance was quantified using Harrell's concordance index (C-index). *Proportional hazards assumption violated: Schoenfeld test $p < 0.05$. Abbreviations: PH, proportional hazards; amp, amplification.

| | IDH-mutant Astrocytoma | | | IDH-mutant Oligodendroglioma | | |
|---|---|---|---|---|---|---|
| | All tumors | Hypermutant | Non-Hypermutant | All tumors | Hypermutant | Non-Hypermutant |
| **Clinical markers** | | | | | | |
| *Demographics* | | | | | | |
| Age of diagnosis (>40 vs ≤40) | 1.64 [1.00-2.71] | 1.71 [0.64-4.55] | **1.90 [1.05-3.45]** | **2.31 [1.25-4.27]** | 1.14 [0.38-3.46] | **3.22 [1.29-8.06]** |
| Sex (male vs female) | 1.30 [0.83-2.03] | 0.67 [0.29-1.53] | **1.80 [1.01-3.21]** | 1.39 [0.74-2.61] | 1.52 [0.48-4.91] | 1.91 [0.73-5.03] |
| *Primary treatment* | | | | | | |
| Surgery type (resection vs biopsy) | *all resections* | *all resections* | *all resections* | 1.34 [0.25-7.23] | *all resections* | 1.06 [0.19-5.90]* |
| Radiotherapy (yes vs no) | 0.66 [0.39-1.10] | 0.95 [0.37-2.47] | 0.59 [0.32-1.11] | 0.75 [0.35-1.59] | 1.58 [0.44-5.64] | 0.79 [0.30-2.08] |
| Alkylating chemotherapy (yes vs no) | **0.49 [0.25-0.94]** | 0.35 [0.11-1.13] | 0.52 [0.22-1.20] | **3.74 [1.35-10.38]** | 1.16 [0.29-4.54] | **5.94 [1.07-32.95]** |
| *Primary pathology* | | | | | | |
| Grade (3 vs other)** | 1.21 [0.55-2.66] | *all grade >2* | 0.98 [0.42-2.28] | **2.83 [1.13-7.06]** | 0.99 [0.24-4.15] | 3.10 [0.86-11.19] |
| Grade (4 vs other)** | **2.27 [1.22-4.22]** | 0.61 [0.17-2.17] | **2.03 [1.05-3.90]** | | | |
| **Molecular markers at recurrence** | | | | | | |
| *Mutations (present vs absent)* | | | | | | |
| *MSH6* | 1.77 [0.91-3.46] | 0.96 [0.42-2.22] | *all wildtype* | **2.96 [1.41-6.19]** | 0.74 [0.23-2.46] | *all wildtype* |
| *NOTCH1* | **1.97 [1.11-3.49]** | 1.12 [0.49-2.57] | 1.29 [0.24-6.95] | **2.15 [1.08-4.28]** | 1.01 [0.33-3.07] | 1.65 [0.56-4.83] |
| *PIK3CA* | 1.24 [0.53-2.91] | 0.78 [0.22-2.72] | 1.33 [0.40-4.40] | **2.54 [1.34-4.80]** | 0.72 [0.19-2.71] | 1.74 [0.66-4.54] |
| *PIK3R1* | 1.45 [0.44-4.74] | 1.05 [0.30-3.70] | <0.001 [0-∞] | 1.16 [0.47-2.84] | 0.59 [0.16-2.20] | 1.21 [0.34-4.24] |
| Hypermutation | **2.09 [1.29-3.37]** | | | **3.71 [1.87-7.35]** | | |
| *Homozygous deletions (HD vs non-HD)* | | | | | | |
| *CDKN2A/B* | **1.72 [1.07-2.74]** | 1.59 [0.64-3.97] | 1.76 [1.02-3.06]* | 15.33 [4.83-48.62]* | *all non-HD* | 21.18 [5.87-76.45]* |
| *PTEN* | *all non-HD* | *all non-HD* | *all non-HD* | *all non-HD* | *all non-HD* | *all non-HD* |
| *ATRX* | 1.22 [0.37-3.99] | **18.49 [1.81-189.40]** | 0.69 [0.13-3.64] | *all non-HD* | *all non-HD* | *all non-HD* |
| *Amplifications (amp vs non-amp)* | | | | | | |
| *CCND2* | 1.81 [0.97-3.40] | 0.54 [0.15-1.91] | **2.90 [1.29-6.47]** | *all non-amp* | *all non-amp* | *all non-amp* |
| *EGFR* | 0.74 [0.22-2.45] | *all non-amp* | 0.86 [0.26-2.86] | *all non-amp* | *all non-amp* | *all non-amp* |
| *PDGFRA* | 0.85 [0.39-1.85] | 0.77 [0.27-2.21] | 0.67 [0.20-2.20] | *all non-amp* | *all non-amp* | *all non-amp* |
| *CDK4/6* | **2.37 [1.29-4.32]** | 2.26 [0.86-5.94] | 2.09 [0.94-4.61] | 1.80 [0.33-9.73] | 4.14 [0.62-27.71] | <0.001 [0-∞] |
| *MET/PTPRZ1* | 1.70 [0.90-3.22] | 1.84 [0.70-4.83] | 1.34 [0.56-3.22] | <0.001 [0-∞] | *all non-amp* | <0.001 [0-∞] |
| *MYC/MYCN* | 1.46 [0.82-2.60]* | **2.68 [1.07-6.70]** | 0.96 [0.44-2.12] | *all non-amp* | *all non-amp* | *all non-amp* |
| *Methylation (methylated vs unmethylated)* | | | | | | |
| *MGMT* promoter | 1.27 [0.46-3.52] | 1.11 [0.19-6.44] | 1.23 [0.35-4.35] | >1.00x$10^5$ [0-∞] | *all methylated* | >1.00x$10^5$ [0-∞] |

**Extended Data Table 5. Univariable Cox proportional hazards models concerning post-recurrence survival of GLASS IDH-mutant glioma patients.**
Hazard ratios with 90% confidence intervals. Significant factors for multivariable testing were defined as all variables with $p < 0.10$ that did not violate the proportional hazards assumption. *Proportional hazards assumption violated: Schoenfeld test $p < 0.05$. **Variable consist of 3 separate levels. Abbreviations: HD, homozygous deletion; amp, amplification.

| | IDH-mutant Astrocytoma | | | IDH-mutant Oligodendroglioma | |
|---|---|---|---|---|---|
| | All tumors | Hypermutant | Non-Hypermutant | All tumors | Non-Hypermutant |

| **Full Multivariable Cox PH Model** | C-index = 0.68 | C-index = 0.61 | C-index = 0.69 | C-index = 0.79 | C-index = 0.72 |
|---|---|---|---|---|---|
| Age of diagnosis (>40 vs ≤40) | | | 1.45 [0.68-3.09] | 2.88 [0.83-9.95] | 2.89 [0.86-9.68] |
| Sex (male vs female) | | | 1.37 [0.66-2.84] | | |
| Alkylating chemotherapy (yes vs no) | **0.34 [0.14-0.83]** | | | 2.82 [0.60-13.29] | 4.53 [0.58-35.70] |
| Grade (3 vs other)* | 1.01 [0.28-3.67] | | 0.95 [0.34-2.62] | 1.52 [0.46-5.02] | |
| Grade (4 vs other)* | 1.11 [0.30-4.16] | | 1.69 [0.75-3.81] | | |
| *MSH6* (mutant vs wildtype) | | | | 0.25 [0.03-1.78] | |
| *NOTCH1* (mutant vs wildtype) | 0.91 [0.28-2.98] | | | 1.79 [0.47-6.89] | |
| *PIK3CA* (mutant vs wildtype) | | | | 0.64 [0.08-5.23] | |
| Hypermutation (present vs absent) | 2.28 [0.73-7.13] | | | **10.63 [0.88-128.30]** | |
| *CDKN2A/B* (HD vs non-HD) | 1.62 [0.58-4.47] | | | | |
| *ATRX* (HD vs non-HD) | | 10.36 [0.59-181.62] | | | |
| *CCND2* (amp vs non-amp) | | | 2.15 [0.80-5.83] | | |
| *CDK4/6* (amp vs non-amp) | **5.93 [1.58-22.28]** | | | | |
| *MYC/MYCN* (amp vs non-amp) | | 2.28 [0.70-7.37] | | | |
| **Stepwise Backward Cox PH Model** | C-index = 0.65 | C-index = 0.55 | C-index = 0.55 | C-index =0.77 | C-index = 0.72 |
| Age of diagnosis (>40 vs ≤40) | | | | 2.38 [0.81-6.98] | 2.89 [0.86-9.68] |
| Alkylating chemotherapy (yes vs no) | **0.40 [0.17-0.92]** | | | 3.11 [0.70-13.82] | 4.53 [0.58-35.70] |
| Hypermutation (present vs absent) | **2.30 [1.08-4.91]** | | | **3.37 [1.27-8.98]** | |
| *ATRX* (HD vs non-HD) | | **18.49 [1.16-295.70]** | | | |
| *CCND2* (amp vs non-amp) | | | **2.88 [1.10-7.55]** | | |
| *CDK4/6* (amp vs non-amp) | **4.92 [1.56-15.51]** | | | | |

**Extended Data Table 6. Multivariable Cox proportional hazards models concerning post-recurrence survival of GLASS IDH-mutant glioma patients.**
Hazard ratios with 95% confidence intervals. Significant factors were defined as all variables with $p < 0.05$ that did not violate the proportional hazards assumption. Model performance was quantified using Harrell's concordance index (C-index). *Variable consist of 3 separate levels. Abbreviations: PH, proportional hazards; HD, homozygous deletion; amp, amplification.
