## Supplementary Figures for "Tumor-initiating genetics and therapy drive divergent molecular evolution in IDH-mutant gliomas"

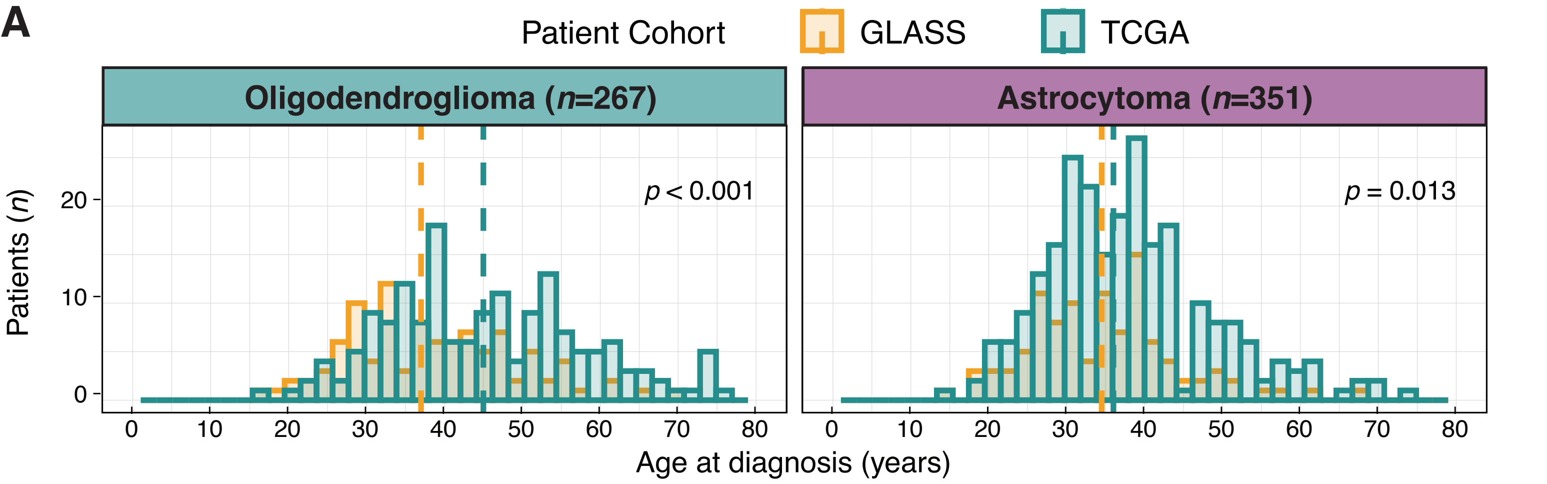


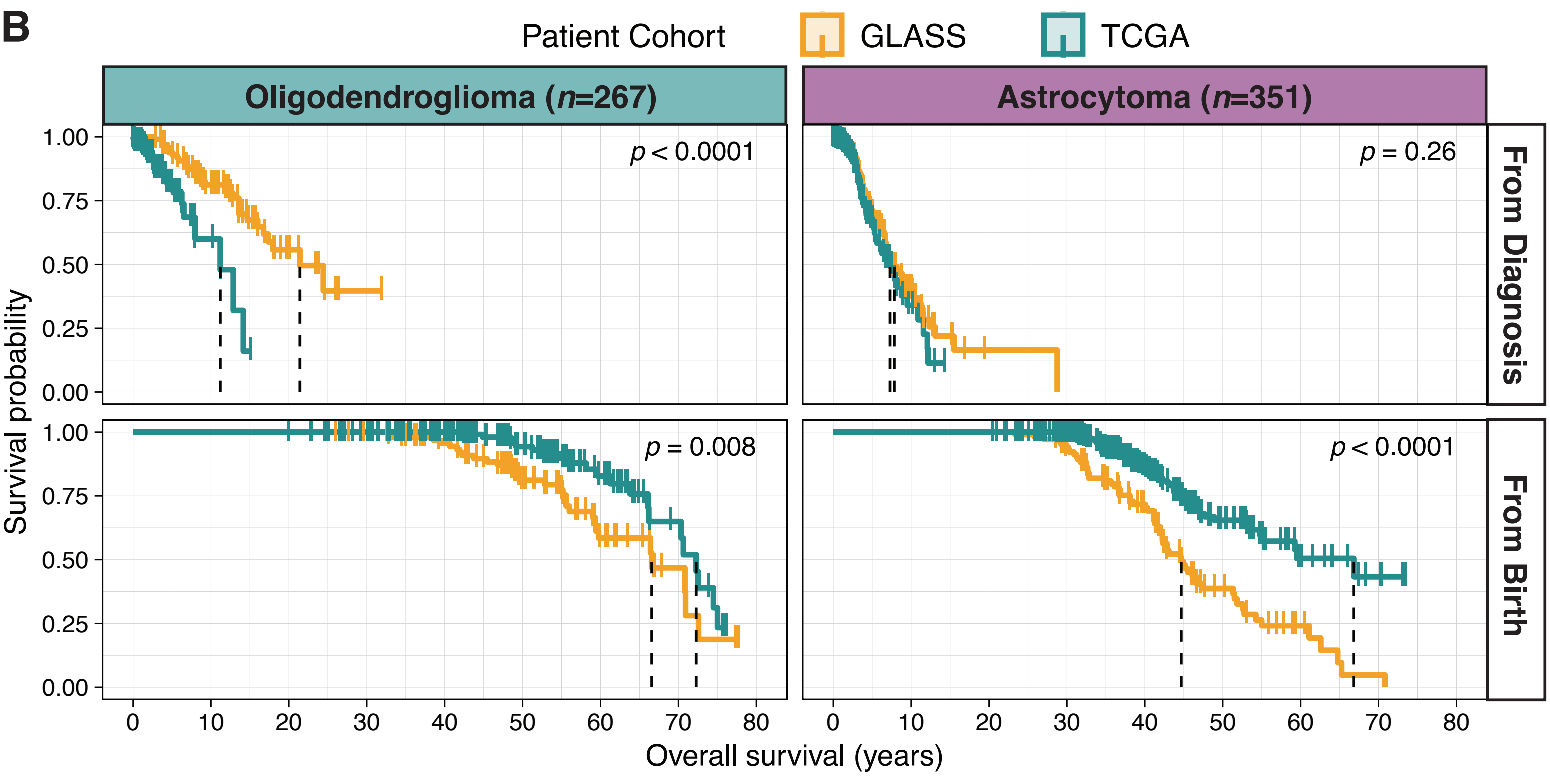


**C**

| | Oligodendroglioma | | Astrocytoma | |
|---|---|---|---|---|
| | **GLASS (*n*=100)** | **TCGA (*n*=167)** | **GLASS (*n*=104)** | **TCGA (*n*=247)** |
| **Events, *n*** | 30 | 21 | 63 | 52 |
| **Overall survival from diagnosis** | | | | |
| Median [95%CI], yrs | 21.4 [17.33-NR] | 11.2 [7.96-NR] | 7.83 [6.67-10.2] | 7.29 [5.62-10.9] |
| HR [95%CI] | 3.68 [1.91-7.11] | | 1.25 [0.85-1.83] | |
| **Overall survival from birth** | | | | |
| Median [95%CI], yrs | 66.6 [59.4-NR] | 72.3 [66.2-NR] | 44.7 [42.2-51.3] | 66.8 [55.2-NR] |
| HR [95%CI] | 0.48 [0.27-0.83] | | 0.36 [0.25-0.53] | |

**Extended Data Figure 1. Early disease onset in GLASS patients is associated with reduced life expectancy compared to those in the TCGA cohort.**
**A.** Age distribution at time of diagnosis of GLASS and TCGA cohorts. Dotted lines represent median age across the respective cohorts. **B and C.** Despite having a longer overall survival from the time of diagnosis for patients with oligodendrogliomas (top Kaplan-Meier panels), GLASS patients with both glioma types had a worse projected overall survival than TCGA patients when measured from birth (bottom Kaplan-Meier panels). Projected overall survival from birth was calculated by adding individually reported overall survival years from diagnosis to age of disease onset. (**B**: Kaplan-Meier estimates, **C**: Tabular summary to emphasize the confidence intervals per group and hazard ratios between groups.) Abbreviations: CI, confidence interval; yrs, years; HR, hazard ratio; NR, not reached.

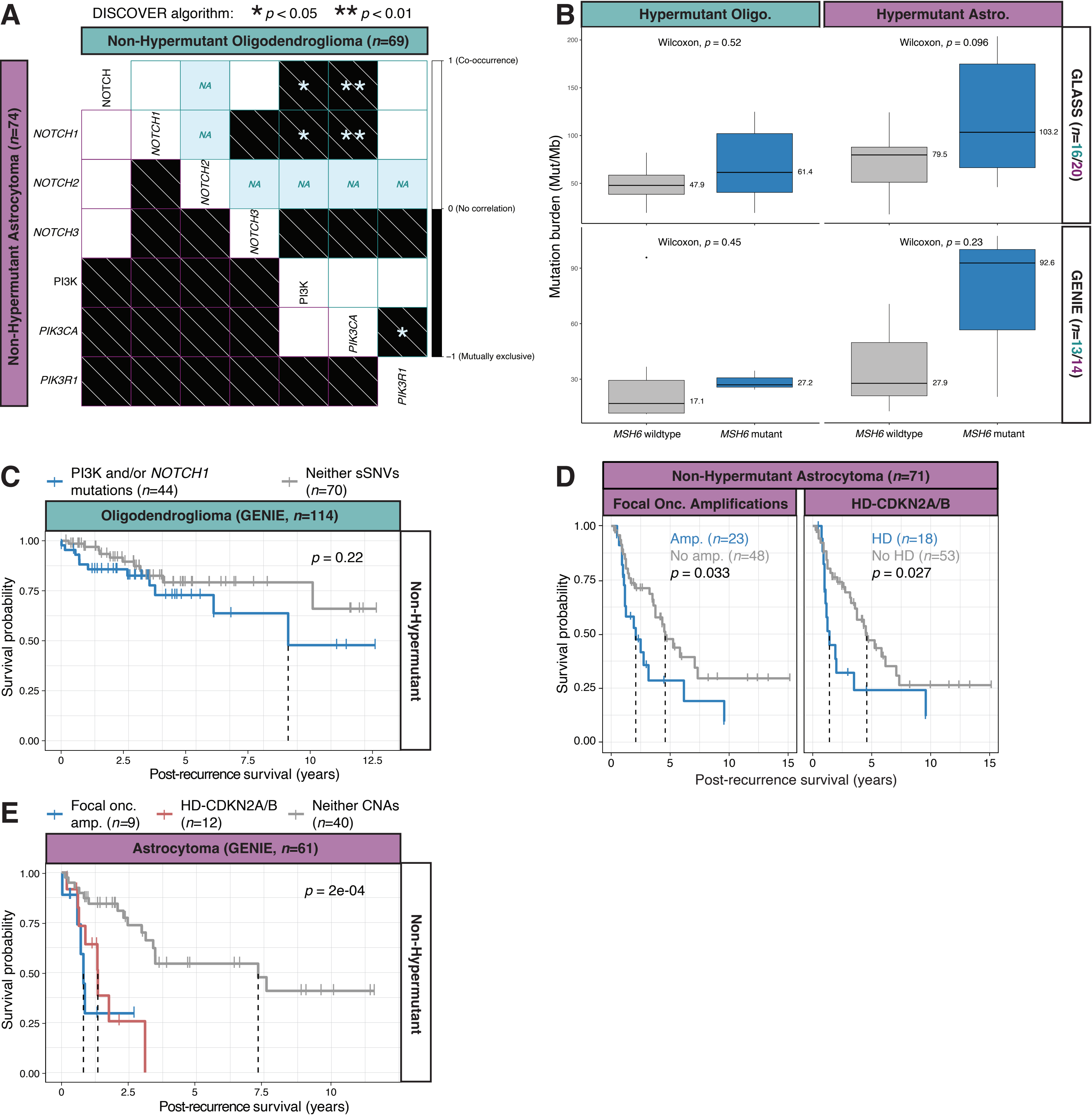


**Extended Data Figure 2. Post-recurrence survival is influenced by specific driver mutations and hypermutation status.**
**A.** Contingency table of mutual exclusivity *p*-values of somatic single-nucleotide variants detected in PI3K and NOTCH genes in non-hypermutant oligodendrogliomas. DISCOVER algorithm results showed mutual exclusivity between PI3K and NOTCH mutations in patients with non-hypermutant oligodendrogliomas but not astrocytomas. **B.** Comparison of mutational burden between *MSH6*-mutant and *MSH6*-wildtype hypermutant oligodendrogliomas and astrocytomas in the GLASS cohort. **C.** Post-recurrence survival in GENIE cohort patients with non-hypermutant oligodendrogliomas stratified for PI3K and *NOTCH1* mutations. **D.** Post-recurrence survival in GLASS cohort patients with non-hypermutant astrocytomas stratified for focal oncogene amplifications and homozygously deleted *CDKN2A/B*. **E.** Post-recurrence survival in GENIE cohort patients with non-hypermutant astrocytomas stratified for focal oncogene amplifications and homozygously deleted *CDKN2A/B*. Abbreviations: Mut, mutations; Mb, megabase; Onc., Oncogene; HD, homozygous deletion; Amp., amplifications; sSNVs, somatic single nucleotide variants; CNAs, copy number alterations.

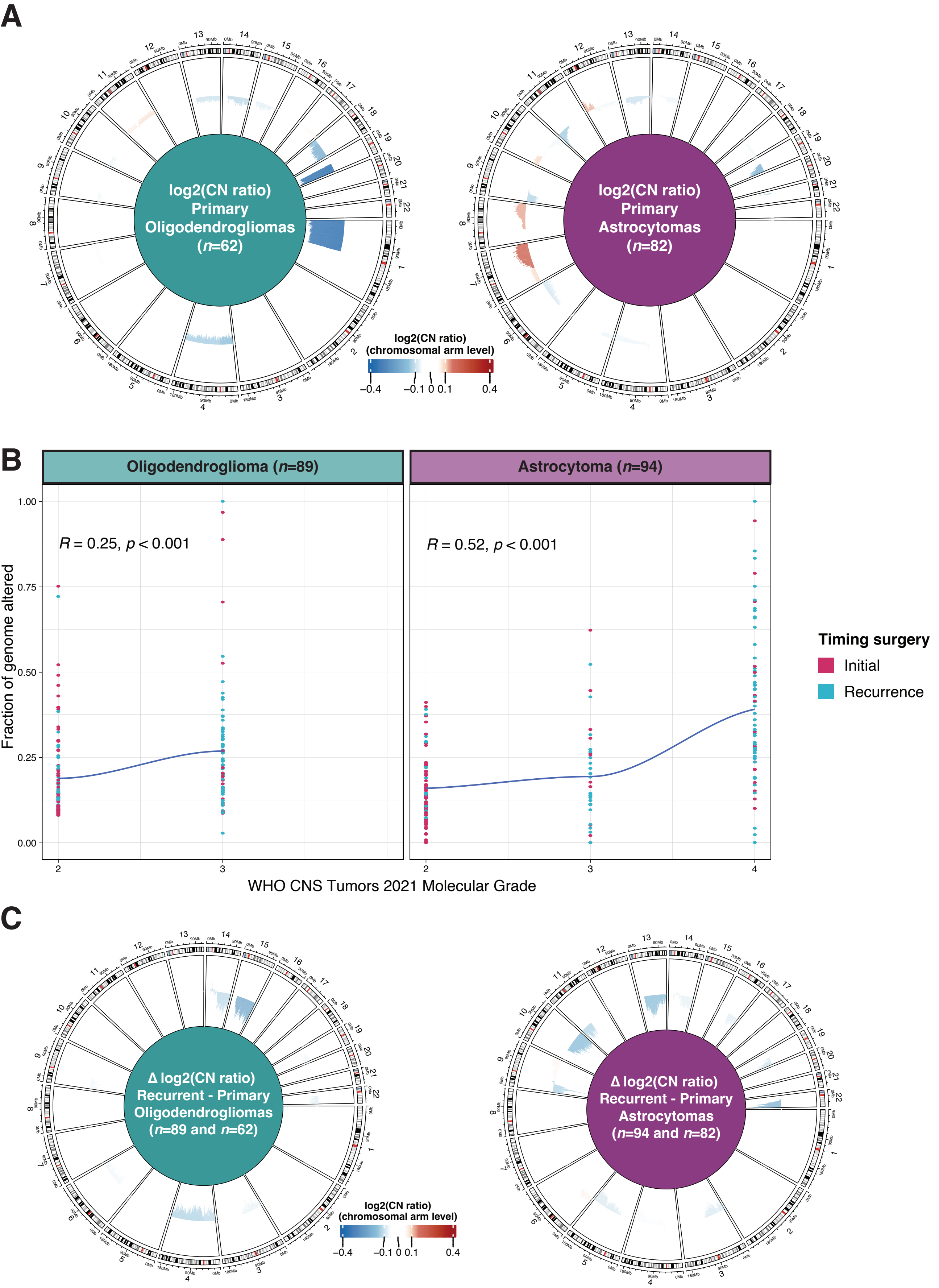


**Extended Data Figure 3. Both glioma types have similar arm-level aneuploidies at primary surgery, but astrocytomas accumulate more over time.**
**A.** Circos plot showing average DNA copy number across primary-only oligodendrogliomas and primary-only astrocytomas. Arm-level aneuploidies (mean log2(CN ratio) ≤ -0.1 or ≥ 0.1) are shown. **B.** Fraction of genome altered by WHO molecular grade. **C.** Comparison of arm-level aneuploidies differences between initial and recurrent oligodendrogliomas and astrocytomas. Arm level events shown are mean log2(CN ratio) ≤ -0.05 or ≥ 0.05. The significant changes in CN ratios from primary to recurrent gliomas indicate ongoing genomic evolution, particularly in astrocytomas. Abbreviations: CN, copy number.

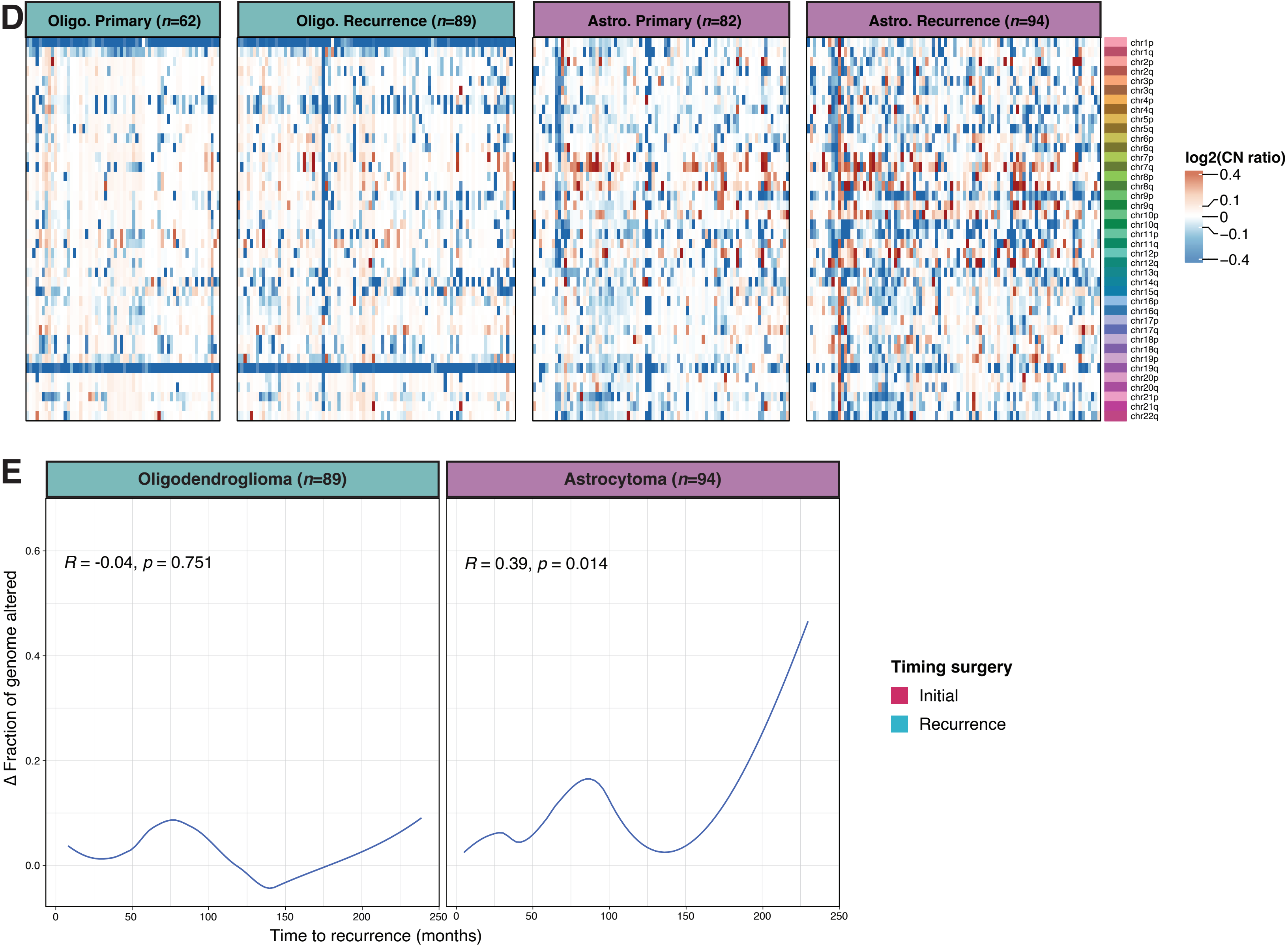


**Extended Data Figure 3 (continued). Both glioma types have similar arm-level aneuploidies at primary surgery, but astrocytomas accumulate more over time.** **D.** Genome-wide DNA copy number profiles of primary-only and matching recurrent oligodendrogliomas and astrocytomas. **E.** Change in fraction of genome altered compared to surgical interval. Abbreviations: Olig., oligodendroglioma; Astro., astrocytoma; CN, copy number.

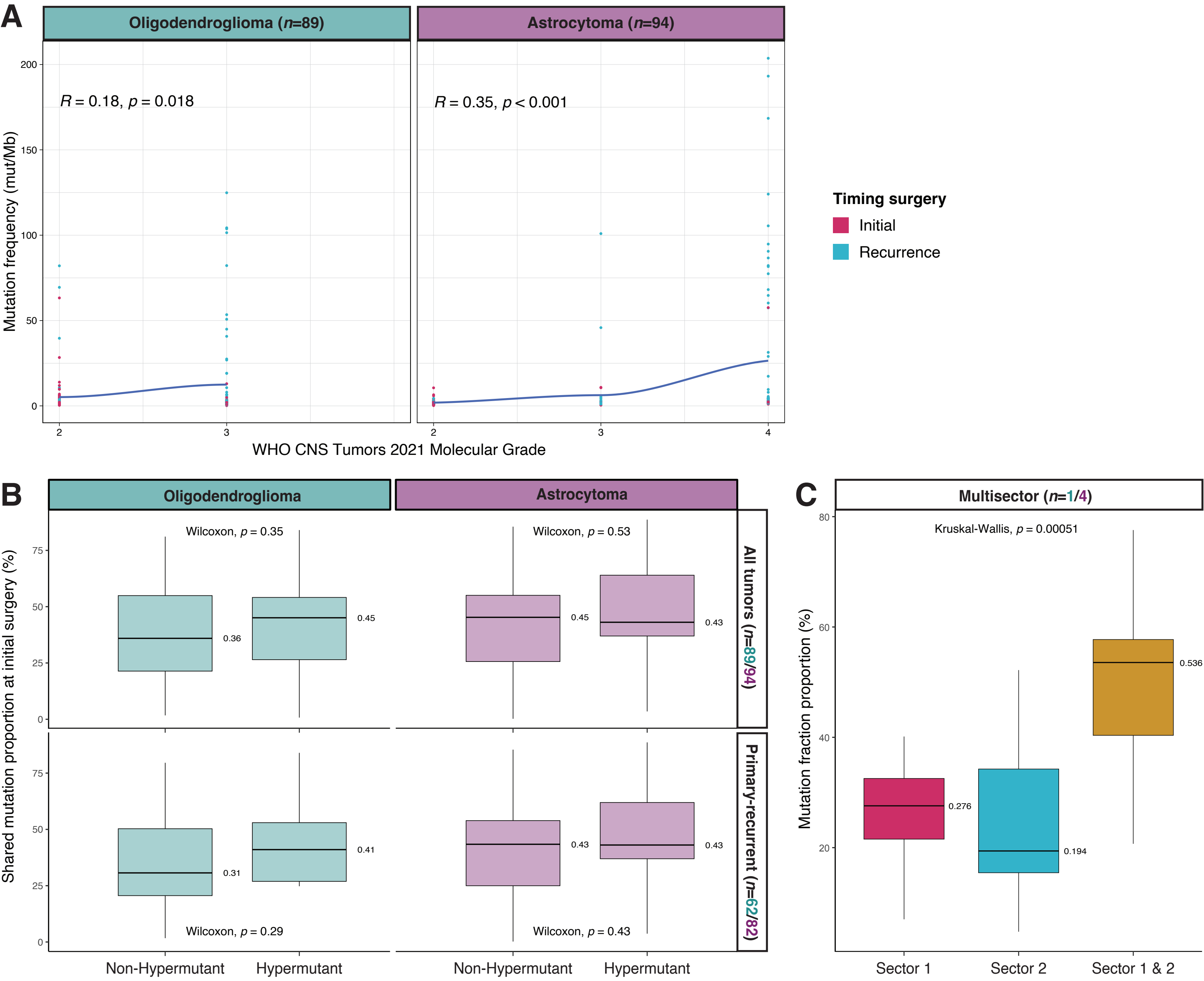


**Extended Data Figure 4. Intratumoral heterogeneity is suggested by variation in mutation distributions across different sectors of the tumor.**
**A.** Comparison of the fraction of mutations that were detected in the initial tumor and retained in the matching recurrent tumor. **B.** Multisector sampling from the same surgery, comparing mutation distributions across different sectors in 1 oligodendroglioma and 4 astrocytomas. Median fraction of mutations detected uniquely in Sector 1 (left), uniquely detected in Sector 2 (middle) or shared between Sectors 1 and 2 (right).

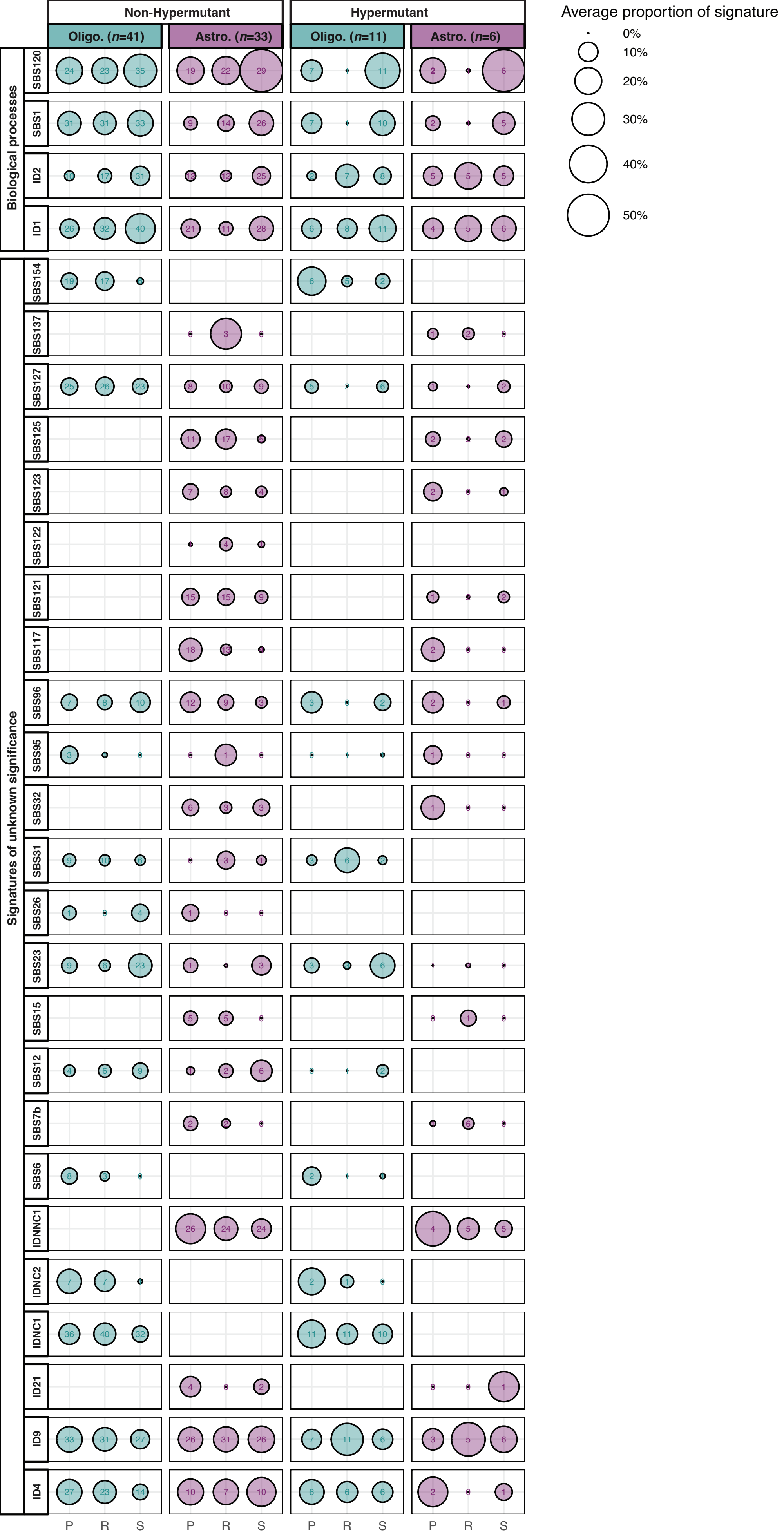


**Extended Data Figure 5. Distinct mutational signatures reveal insights into the etiology and evolution of gliomas.**
Comparison of SBS and ID signatures using the average percentage of alterations attributed to these signatures (circle size) and the number of patients with any alteration attributable to these signatures (number in circle) per glioma type, hypermutation status, treatment regimen, and timing of the signatures. Abbreviations: Olig., Oligodendroglioma; Astro., Astrocytoma; SBS, single-base substitutions; ID, small insertions and deletions; P, primary; R, recurrence; S, shared.
